# Plasmid-mediated metronidazole resistance in *Clostridioides difficile*

**DOI:** 10.1101/643775

**Authors:** Ilse M. Boekhoud, Bastian V. H. Hornung, Eloisa Sevilla, Céline Harmanus, Ingrid M. J. G. Bos-Sanders, Elisabeth M. Terveer, Rosa Bolea, Jeroen Corver, Ed J. Kuijper, Wiep Klaas Smits

## Abstract

**Background:** Metronidazole is used to treat mild- to moderate *Clostridioides difficile* infections (CDI). No clear mechanism for metronidazole resistance has been described for *C. difficile*. A patient treated in the Leiden University Medical Center suffered from recurrent CDI caused by a PCR ribotype (RT) 020 strain which developed resistance to metronidazole (MIC = 8 mg/L). Resistance is also seen in animal isolates, predominantly of RT010.

**Methods:** Six metronidazole susceptible and 12 metronidazole resistant isolates from human and animal origin, including the patient isolates, were analyzed by whole genome sequence (WGS) analysis. 585 susceptible and resistant isolates collected in various international studies were tested for the presence of plasmid by PCR. Plasmid copy number was determined by quantitative PCR.

**Findings:** Stable metronidazole resistance correlated with the presence of a 7kb plasmid, pCD-METRO. pCD-METRO was not detected in 562 susceptible isolates, but was found in toxigenic and non-toxigenic metronidazole resistant strains from multiple countries (n=22). The introduction of a pCD-METRO-derived vector into a susceptible strain led to a ∼25 fold increase in the metronidazole MIC. The pCD-METRO replicon sustained a plasmid copy number of ∼30, which is higher than currently known replicons for *C. difficile*.

**Interpretation:** We describe the first plasmid-mediated resistance to a clinically relevant antibiotic in *C. difficile*. pCD-METRO is an internationally disseminated plasmid capable of conferring metronidazole resistance in *C. difficile*, including epidemic ribotypes. Our finding that pCD-METRO may be mobilizable can impact diagnostics and treatment of CDI.

**Funding:** Netherlands Organisation for Scientific Research; Netherlands Center for One Health; European Center for Disease Prevention and Control

**Research in context:** *Evidence before this study:* On October 19, 2017, a PubMed search was performed with the terms ‘metronidazole resistance’ and ‘clostridium OR clostridioides’, without language restrictions. A single relevant paper was found describing a strain displaying stable metronidazole resistance not obtained by serial passaging, but no mechanism was identified in this study. On the same day, a PubMed search using terms ‘plasmid’ and ‘resistance’ and ‘clostridium difficile OR clostridioides difficile’ did not yield relevant literature on plasmid-mediated resistance in *C. difficile*.

*Added value of this study:* This study is the first report of plasmid-mediated resistance in *C. difficile*, and more generally, the first to ascribe a clinically relevant function to a *C. difficile* plasmid. Specifically, we report the sequence and annotation of the plasmid pCD-METRO and show that it confers stable resistance to metronidazole, is detected in both toxigenic and non-toxigenic strains of human and animal origin (including epidemic types), is internationally disseminated, is maintained at a higher copy number than characterized *C. difficile* plasmids and can be acquired horizontally.

*Implications of all the available evidence:* Metronidazole is widely used as a treatment for mild-to-moderate CDI, though treatment failure occurs in up to ∼30 % of patients. Our data show that carriage of pCD-METRO results in stable metronidazole resistance in *C. difficile* and suggest that pCD-METRO is mobilizable from an as-of-yet unknown bacterium. Our findings warrant a further investigation into the role of this plasmid in metronidazole treatment failure and the influence of metronidazole use on the international dissemination of pCD-METRO. It also offers an opportunity to improve treatment success and reduce the dissemination of antimicrobial resistance by screening *C. difficile* isolates or donor fecal material prior to fecal microbiota transplant.

## Introduction

*Clostridioides difficile* (*Clostridium difficile*) is a gram-positive obligate anaerobe capable of causing *Clostridium difficile* Infection (CDI) upon disruption of the normal intestinal flora.^1^ Although it is one of the major causes of nosocomial infectious diarrhea, community-acquired CDI is becoming more frequent.^2, 3^ CDI infection poses a significant economic burden with an estimated cost at €3 billion per year in the European Union and impairs the quality of life in infected individuals.^4, 5^ The incidence of CDI has increased over the last two decades with outbreaks caused by epidemic types such as PCR ribotype (RT) 027 (NAP1/BI).^6^ CDI is not restricted to this type, however, as infections caused by RT001, RT002, RT014/020 and RT078 are frequently reported in both Europe and the United States.^7, 8^ Metronidazole is frequently used for the treatment of mild-to-moderate infections and vancomycin for severe infections, though vancomycin is increasingly indicated as a general first-line treatment ^9, 10^ Fidaxomicin has recently also been approved for CDI treatment, but its use is limited by high costs.^11^ Fecal Microbiota Transplantation (FMT) is effective at treating recurrent CDI (rCDI) that is refractory to antimicrobial therapy. ^12^ Reduced susceptibility and resistance to clinically used antimicrobials, including metronidazole, has been reported and this, combined with the intrinsic multiple drug-resistant nature of *C. difficile,* stresses the importance for the development of new effective treatment modalities.^8^

Routine antimicrobial susceptibility testing is generally not performed for *C. difficile* and consequently, reports of resistance to metronidazole are rare.^13–15^ Longitudinal surveillance in Europe found that 0·2% of clinical isolates investigated were resistant to metronidazole, ^14^ but reported rates from other studies vary from 0-18·3%.^16–19^ These differences may reflect geographic distributions in resistant strains, or differences in testing methodology and breakpoints used.^20, 21^ Moreover, metronidazole resistance can be unstable, inducible and heterogeneous.^22^ Finally, metronidazole resistance appears to be more frequent in non-toxigenic strains such as those belonging to RT010, which have a 7-9 fold increase in Minimal Inhibitory Concentration (MIC) values compared to RT001, RT027 and RT078. ^16, 21^

Metronidazole is a 5-nitroimidazole drug that upon intracellular reductive activation induces cellular damage through nitrosoradicals.^22^ Mechanisms associated with metronidazole resistance described in other organisms include the presence of 5-nitroimidazole reductases (*nim* genes), altered pyruvate-ferredoxin oxidoreductase (PFOR) activity and adaptations to (oxidative) stress.^22^ The knowledge on resistance mechanisms in *C. difficile* is very limited, but may involve modulation of core metabolic and stress pathways as well.^23^

Here, we present a case of a patient with rCDI due to an initially metronidazole susceptible (MTZ^S^) RT020 strain which developed resistance to metronidazole over time. We analyzed the genome sequences of these toxigenic MTZ^S^ and metronidazole resistant (MTZ^R^) strains, together with 4 MTZ^S^ and 8 MTZ^R^ non-toxigenic RT010 strains. We identified pCD-METRO, a 7-kb plasmid conferring metronidazole resistance. This plasmid is internationally disseminated and also occurs in epidemic types. This is the first report of a clinically relevant phenotype associated with plasmid-carriage in *C. difficile*.

## Methods

### Strains

The 18 strains sequenced as part of this study were isolated from a single patient at the Leiden University Medical Center (LUMC) or derived from the collection of the Dutch National Reference Laboratory (NRL) for *C. difficile*, which is hosted at the LUMC. Informed consent was given for the use of the patient samples for research purposes. Other clinical isolates (n=567) were obtained through the NRL and partners in the *C. difficile* typing network of the European Centre for Disease Prevention and Control (ECDC), or were previously collected as part of the ECDIS study and the Tolevamer and MODIFY I+II clinical trials.^24–26^

### Whole genome sequencing and analysis

DNA was extracted from 9mL of stationary growth phase cultures grown in Brain-Heart Infusion (BHI, Oxoid) broth using a QIAsymphony (Qiagen, The Netherlands) with the QIAsymphony DSP Virus/Pathogen Midi Kit according to the manufacturer’s instructions. All samples were sequenced on an Illumina HiSeq4000 platform with read length 150bp in paired-end mode, followed by assembly, annotation, SNP calling and plasmid analysis using in-house pipelines (described in detail in the appendix).

### Data accessibility

All sequence data generated in this study has been uploaded to the European Nucleotide Archive under project PRJEB24167 with accession numbers ERR2232520- ERR2232537. The genome assembly for IB136, including the annotated sequence of pCD-METRO, can be found under accession number ERZ807316.

### Antimicrobial susceptibility testing and ribotyping

All strains were characterized by standardized PCR ribotyping and tested for metronidazole resistance by agar dilution according to Clinical & Laboratory Standards Institute guidelines using the EUCAST epidemiological cut-off of 2mg/L.^27–29^ Details of all strains and their characteristics are available in table 1 and supplemental table 1 (appendix). For epsilometer tests (E-test; BioMerieux), bacterial suspensions corresponding to 1·0 McFarland turbidity were applied on BHI agar supplemented with 0·5% yeast extract (Sigma-Aldrich) and *Clostridium difficile* Selective Supplement (CDSS, Oxoid). MIC values were read after 48 hours of incubation.

**Table 1:**
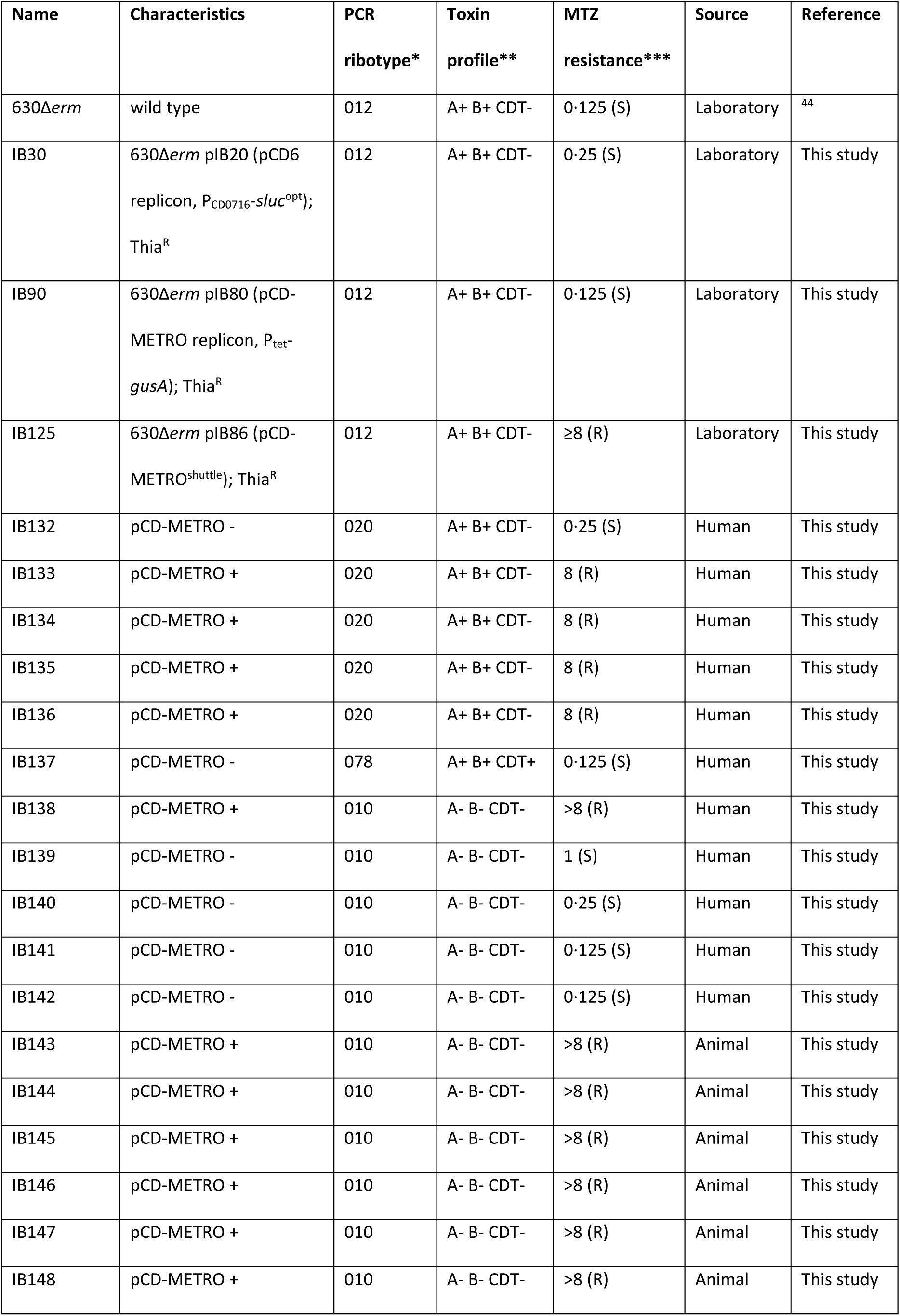
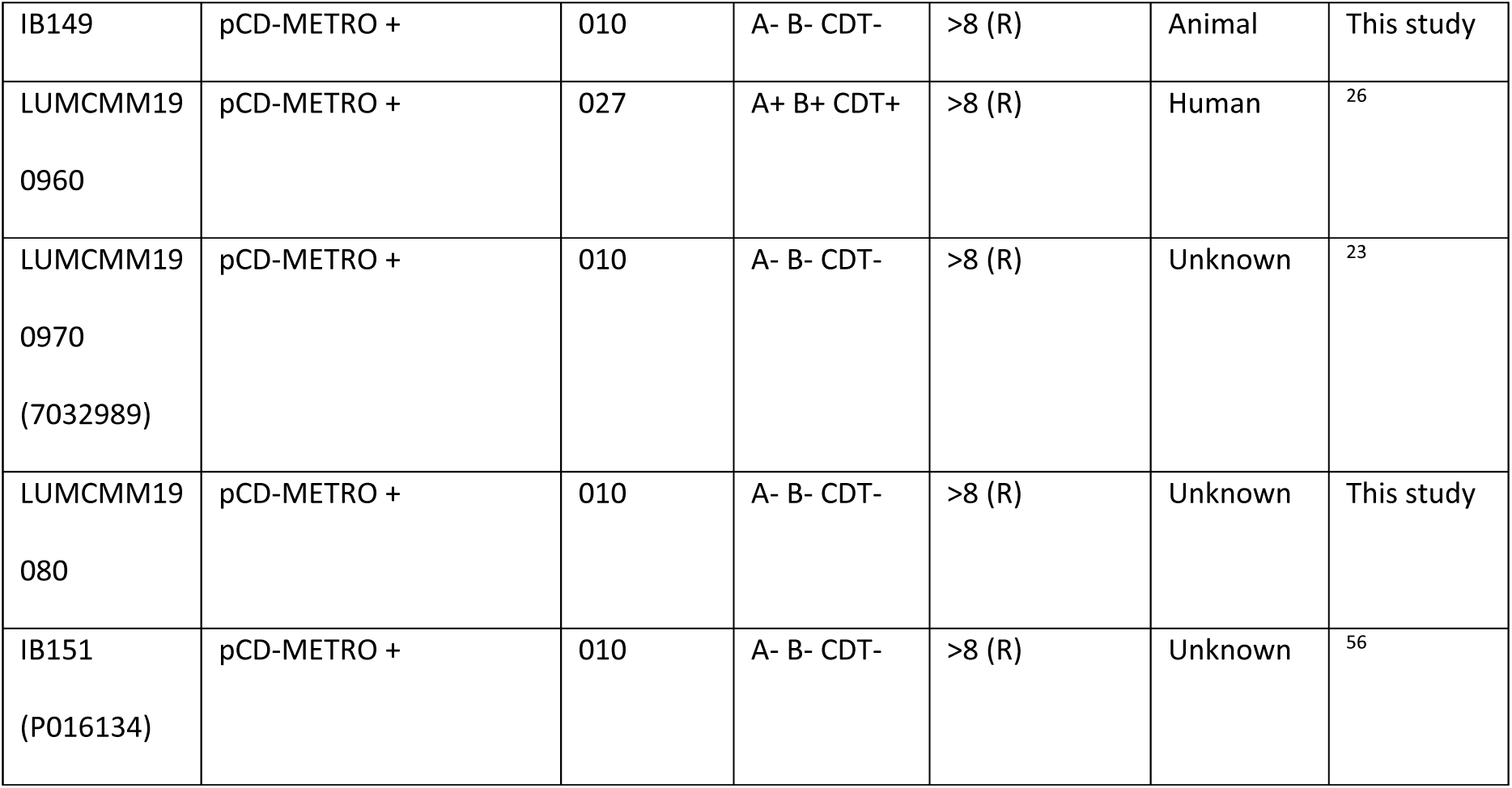
Strains described in this study. Listed are strains mentioned in the main body of the manuscript. For a complete overview of all strains used, see supplemental table 1 (appendix). * PCR ribotype determined at the LUMC using capillary PCR ribotyping ** toxin profile determined by multiplex PCR. *** Metronidazole MIC values in mg/L as determined by agar dilution conform CLSI guidelines. S = susceptible (MIC<2 mg/L), R = resistant (MIC>2 mg/L).

| Name | Characteristics | PCR<br>ribotype* | Toxin<br>profile** | MTZ<br>resistance*** | Source | Reference |
| --- | --- | --- | --- | --- | --- | --- |
| 630Δerm | wild type | 012 | A+ B+ CDT- | 0.125 (S) | Laboratory | <sup>44</sup> |
| IB30 | 630Δerm pIB20 (pCD6 replicon, P<sub>CD0716</sub>-*sluc*<sup>opt</sup>); Thia<sup>R</sup> | 012 | A+ B+ CDT- | 0.25 (S) | Laboratory | This study |
| IB90 | 630Δerm pIB80 (pCD-METRO replicon, P<sub>tet</sub>-*gusA*); Thia<sup>R</sup> | 012 | A+ B+ CDT- | 0.125 (S) | Laboratory | This study |
| IB125 | 630Δerm pIB86 (pCD-METRO<sup>shuttle</sup>); Thia<sup>R</sup> | 012 | A+ B+ CDT- | ≥8 (R) | Laboratory | This study |
| IB132 | pCD-METRO - | 020 | A+ B+ CDT- | 0.25 (S) | Human | This study |
| IB133 | pCD-METRO + | 020 | A+ B+ CDT- | 8 (R) | Human | This study |
| IB134 | pCD-METRO + | 020 | A+ B+ CDT- | 8 (R) | Human | This study |
| IB135 | pCD-METRO + | 020 | A+ B+ CDT- | 8 (R) | Human | This study |
| IB136 | pCD-METRO + | 020 | A+ B+ CDT- | 8 (R) | Human | This study |
| IB137 | pCD-METRO - | 078 | A+ B+ CDT+ | 0.125 (S) | Human | This study |
| IB138 | pCD-METRO + | 010 | A- B- CDT- | >8 (R) | Human | This study |
| IB139 | pCD-METRO - | 010 | A- B- CDT- | 1 (S) | Human | This study |
| IB140 | pCD-METRO - | 010 | A- B- CDT- | 0.25 (S) | Human | This study |
| IB141 | pCD-METRO - | 010 | A- B- CDT- | 0.125 (S) | Human | This study |
| IB142 | pCD-METRO - | 010 | A- B- CDT- | 0.125 (S) | Human | This study |
| IB143 | pCD-METRO + | 010 | A- B- CDT- | >8 (R) | Animal | This study |
| IB144 | pCD-METRO + | 010 | A- B- CDT- | >8 (R) | Animal | This study |
| IB145 | pCD-METRO + | 010 | A- B- CDT- | >8 (R) | Animal | This study |
| IB146 | pCD-METRO + | 010 | A- B- CDT- | >8 (R) | Animal | This study |
| IB147 | pCD-METRO + | 010 | A- B- CDT- | >8 (R) | Animal | This study |
| IB148 | pCD-METRO + | 010 | A- B- CDT- | >8 (R) | Animal | This study |
| IB149 | pCD-METRO + | 010 | A- B- CDT- | >8 (R) | Animal | This study |
| LUMCMM19<br>0960 | pCD-METRO + | 027 | A+ B+ CDT+ | >8 (R) | Human | <sup>26</sup> |
| LUMCMM19<br>0970<br>(7032989) | pCD-METRO + | 010 | A- B- CDT- | >8 (R) | Unknown | <sup>23</sup> |
| LUMCMM19<br>080 | pCD-METRO + | 010 | A- B- CDT- | >8 (R) | Unknown | This study |
| IB151<br>(P016134) | pCD-METRO + | 010 | A- B- CDT- | >8 (R) | Unknown | <sup>56</sup> |

### Molecular biology techniques

*Escherichia coli* was cultured aerobically at 37°C in Luria-Bertani (LB) broth, supplemented with 20 µg/ml chloramphenicol and 50 µg/ml kanamycin when appropriate. *C. difficile* was cultured in BHI supplemented with 0·5% yeast extract, CDSS and 20 µg/ml thiamphenicol when appropriate, in a Don Whitley VA-1000 workstation (10% CO_2_, 10% H_2_ and 80% N_2_ atmosphere).

Plasmids and oligonucleotides are listed in supplemental tables 2 and 3 (appendix), respectively. Plasmid pIB80 was constructed by ATUM (Newark, CA) and contains a pCD-METRO derived fragment inserted in between the KpnI and NcoI sites of pRPF185.^30^ pIB86 was constructed using Gibson assembly using HaeIII-linearized pCD-METRO and a fragment from pRPF185. This fragment was obtained by PCR, and contained the requirements for maintenance in, and transfer from, *E. coli*. A detailed description of these constructions is available in the appendix. Plasmids were maintained in *E. coli* DH5a or MDS42 and verified by Sanger sequencing.^31, 32^

Transfer of plasmids from *E. coli* CA434 to *C. difficile* strains was done using standard methods.^33^ Routine DNA extractions were performed using the Nucleospin Plasmid Easypure (Macherey-Nagel) and DNeasy Blood and Tissue (Qiagen) kits after incubating the cells in an enzymatic lysis buffer according to instructions of the manufacturers.

### Plasmid copy number determination

Real time quantitative PCR (qPCR) experiments were performed essentially as described.^34^ In short, total DNA was isolated after 17h of growth using a phenol-chloroform extraction protocol and diluted to a concentration of 10 ng/µl. 4 µl of the diluted DNA sample was added to 6 µl of a mixture containing SYBR Green Supermix (Bio-Rad) and gene-specific primers (0·4 µM total) for a total volume of 10 µl per well. Gene specific primers used were targeting *rpoB* (chromosome) and *catR* (plasmid) and copy number was calculated using the ΔC_T_ method. Statistical significance was calculated using two-way analysis of variance (ANOVA) and Tukey’s test for multiple comparisons (Prism 8, GraphPad).

## Results

### In-patient development of a metronidazole resistant toxigenic strain

A 54 year old kidney-pancreas transplant patient with a medical history of Type I diabetes mellitus, vascular disease and a double lower leg amputation was on hemodialysis when developing diarrhea. The patient was subsequently diagnosed with CDI and a toxigenic metronidazole sensitive (MIC=0·25 mg/L) RT020 strain was isolated from the fecal material of the patient. Treatment with metronidazole was started, leading to initial resolution of the symptoms (figure 1). Two more episodes of CDI occurred during which the patient was treated primarily with vancomycin prior to an FMT provided by the Netherlands Donor Faeces Bank. At the start of the second episode a MTZ^S^ RT020 strain was once more isolated.

**Figure 1:**
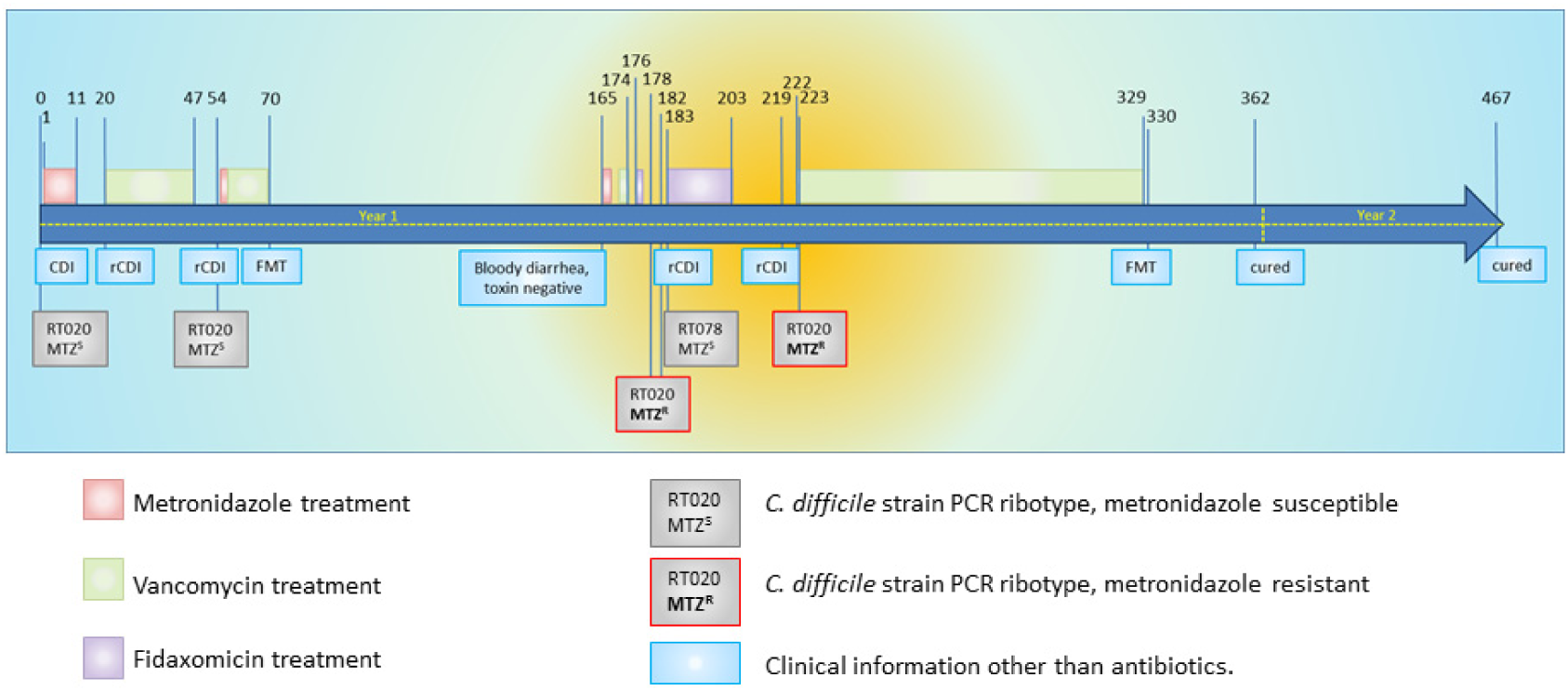
Timeline of the course of the antibiotic treatment and rCDI in the patient. Dates and timepoints on which treatment was initiated or stopped and *C. difficile* isolates were recovered are indicated above the timeline. (r)CDI was diagnosed when both a toxin enzyme-immune assay and nucleic acid amplification test were positive for *C. difficile* in combination with a physician’s assessment of symptoms consistent with CDI. Yellow highlighting indicates the time where pCD-METRO positive MTZ^R^ *C. difficile* was isolated.

Three months after the first FMT, the patient once again developed bloody diarrhea and two more episodes of rCDI were diagnosed which were treated with a vancomycin and a fidaxomicin regime. At two instances, RT020 strains were again isolated from the fecal material of the patient. Strikingly, these two clinical isolates were now phenotypically resistant to metronidazole (MIC=8 mg/L as determined by agar dilution). Ultimately the patient was cured by a second FMT.

We hypothesized that the rCDI episodes were due to clonal RT020 strains that persisted despite antimicrobial therapy and a FMT. Clonal MTZ^S^ and MTZ^R^ strains would allow us to determine the underlying genetic changes that resulted in metronidazole resistance. To determine the relatedness between these RT020 isolates whole genome sequencing (WGS) was performed (table 1). We also included a non-related RT078 strain isolated from the same patient and 4 MTZ^S^ and 8 MTZ^R^ RT010 strains from our laboratory collection (supplemental table 1, appendix) to perform single nucleotide polymorphism (SNP) analyses. Strains were considered resistant to metronidazole with MIC values >2 mg/L according to the EUCAST epidemiological cut-off value.^27^ All strains resistant to metronidazole (n=12) showed cross-resistance to the nitroimidazole drug tinidazole (data not shown).

Assembly of the MTZ^R^ RT020 strain IB136 (see appendix) resulted in a genome of 4166362 bp with 57 contigs, and an average G+C-content of 28·5% (N50= 263391 bp, mapping rate 98·97%). A BLAST comparison between this genome and the NCBI nt database showed that the genome is closest to the genome of strain LEM1.^35^ As expected, 5/6 strains isolated from the patient (all RT020) showed 100% identity over the majority of all contigs, suggesting they are highly similar. The sixth strain, IB137 (RT078), was a clear outlier and was identified as being closest to strain M120, the RT078 reference strain, consistent with the different ribotype assignment.^36^

### Metronidazole resistance does not correlate with a SNP across multiple isolates

Previous studies analysing the mechanism behind metronidazole resistance in *C. difficile* only studied one single isolate.^23, 37^ We performed a core genome SNP analysis on all WGS obtained for this study (n=18; 6 MTZ^S^, 12 MTZ^R^; see table 1), comparing MTZ^S^ and MTZ^R^ strains within and between the different PCR ribotypes (RT010, RT020 and RT078).

The evolutionary rate of *C. difficile* has been estimated at 0-2 SNPs/genome/year but might vary based on intrinsic (strain type) and extrinsic (selective pressure) factors.^38^ Our analysis identified a single SNP in MTZ^R^ RT020, compared to the MTZ^S^ RT020 strains derived from the same patient, conclusively demonstrating that these strains are clonal. This implies the MTZ^S^ RT020 strain acquired metronidazole resistance. In contrast, between the MTZ^S^- and MTZ^R^ RT010 isolates (which come from diverse human and animal sources) 457 SNPs were detected. Moreover, RT010 and RT020 were separated by >25.000 SNPs.

The SNP identified in the RT020 strains discriminating the MTZ^S^ from the MTZ^R^ isolates is located in a conserved putative cobalt transporter (CbiN, IPR003705). However, the SNP is not observed in the MTZ^R^ RT010 strains. Thus, metronidazole resistance is either multifactorial or not contained within the core genome. We did not investigate the contribution of this SNP to metronidazole resistance further.

### MTZ^R^ *C. difficile* strains contain a 7-kb plasmid

Next, we investigated extrachromosomal elements (ECEs), which can include plasmids. Although plasmids containing antimicrobial resistance determinants have been described in gram-positive organisms, they appear to be more common in gram-negatives.^39^ Plasmids in *C. difficile* are known to exist, but no phenotypic consequences of plasmid carriage have been described to date.^40^ The investigation of the pan-genome of all sequenced strains, including a prediction of ECEs predicted by an in-house pipeline similar to PLACNET (appendix),^41^ showed a single contig that was present in all MTZ^R^ strains (4·6% - 19·27% of reads mapped, with a minimum of 479.497), but absent from MTZ^S^ strains, of both RT010 and RT020 (0% of reads mapped with a maximum 327 reads). Circularization based on terminal repeats yielded a putative plasmid of 7056bp with a G+C-content of 41·6% (figure 2a). Correct assembly was confirmed by PCR (figure 2b) and Sanger sequencing (data not shown).

**Figure 2:**
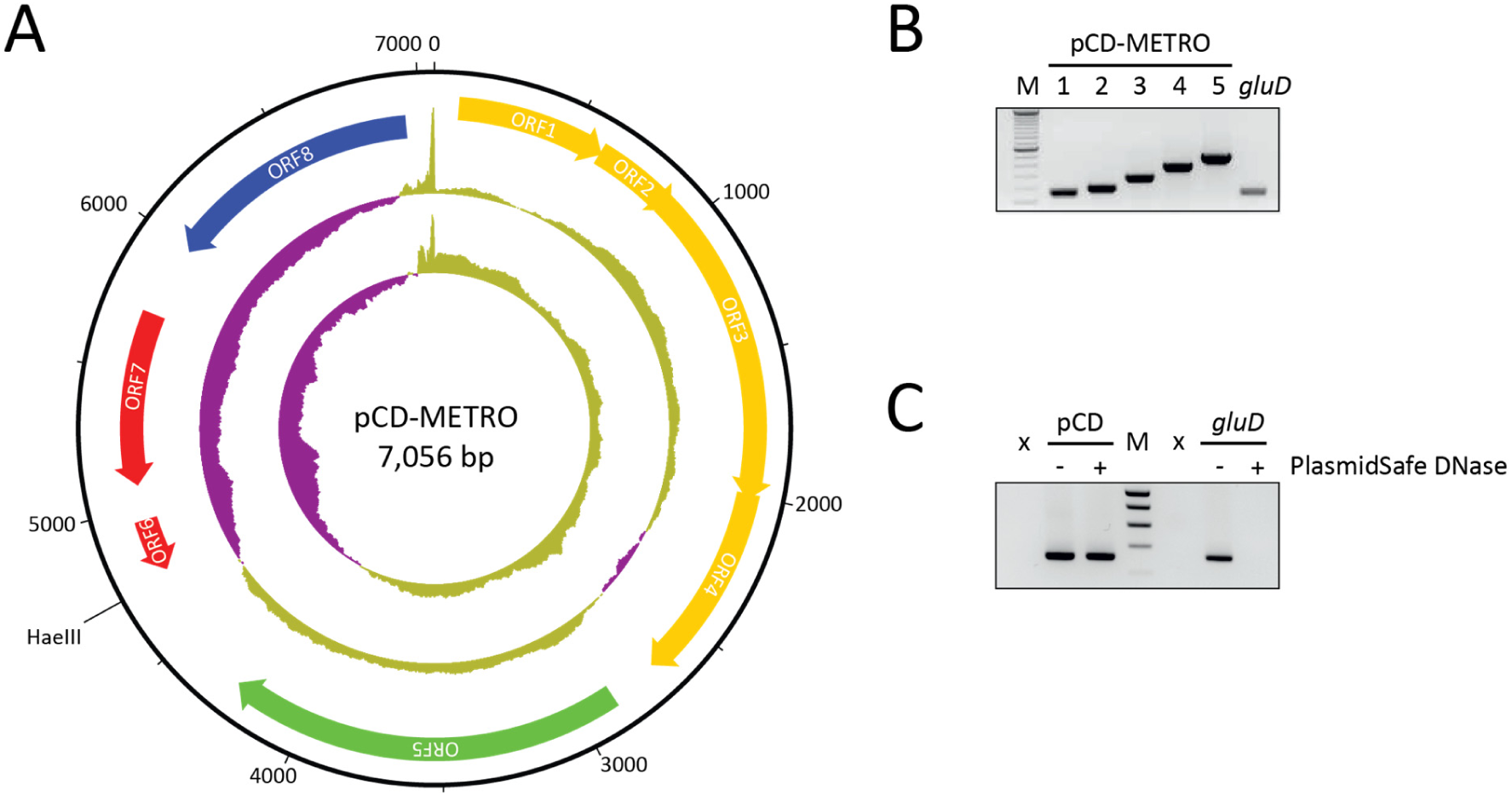
pCD-METRO. (A) Structure of plasmid pCD-METRO and its ORFs. The two innermost circles represent GC content (outer circle) and GC skew (innermost circle) (both step size 5nt and window size 500nt;, above average in yellow, below average in purple). The unique HaeIII site used to construct pCD-METRO^shuttle^ (see methods) is indicated. (B) Gene specific PCR products amplifying regions of ORFs 6 (lane 1+2), ORF5 (lane 3), ORF7 (lane 4) and ORF4 (lane 5), and a chromosomal locus (*gluD*) (C) The product of plasmid specific amplification (targeting ORF6) or chromosomal specific amplification (*gluD*) before and after PlasmidSafe DNase treatment.

To confirm the circular nature of the contig, total DNA isolated from the MTZ^R^ RT010 strain IB138 before and after PlasmidSafe DNase (PSD, Epicentre)^40^ treatment was analyzed by PCR using primers specific for chromosomal DNA (*gluD*) and the putative plasmid (figure 2c). A positive signal for *gluD* was only observed in samples that had not been treated with PSD, demonstrating that PSD treatment degrades chromosomal DNA to below the detection limit of the PCR. By contrast, a signal specific for the putative plasmid was visible both before and after PSD treatment. Consequently, we conclude that our whole genome sequence identified a legitimate 7-kb plasmid.

A total of 8 open reading frames (ORFs) were annotated on the plasmid (figure 2a). ORF1-5 encode a hypothetical protein (ORF1), a MobC-like relaxase/Arc-type ribbon-helix-helix (ORF2; PF05713), a MobA/VirD2 family endonuclease relaxase protein (ORF3; PF03432), a hypothetical protein with a MutS2 signature (ORF4), and a predicted replication protein (ORF5), respectively. ORF6 is a small ORF that is likely a pseudogene, and the remaining ORFs encode a metallohydrolase/oxidoreductase protein (ORF7; IPR001279) and a Tn*5*-like transposase gene (ORF8; PF13701). Intriguingly, ORF6 showed homology on the protein level to the 5-nitroimidazole reductase (*nim*) gene *nimB* (33% identity, 54% positives over 61 amino acids) described in *Bacteroides fragilis* (CAA50578.1) and found in both metronidazole resistant- and susceptible isolates of anaerobic gram-positive cocci.^42, 43^ The ORF lacks the region encoding the N-terminal part of the Nim protein, and the Phyre2-predicted protein structure shows it lacks the catalytic site residues (data not shown). Of note, the plasmid sequences from all strains are highly similar. Compared to the plasmid of strain IB136, only strains IB143, IB144 and IB145 contained a single SNP resulting in a Y314S mutation within the Tn*5*-like transposase ORF.

Together, these results show that all of the MTZ^R^ strains, but none of the MTZ^S^ strains, sequenced in this study contain a plasmid hereafter referred to as pCD-METRO (for **p**lasmid from ***C***. ***d**ifficile* associated with **metro**nidazole resistance).

### pCD-METRO is found in metronidazole resistant strains from different countries

Very few clinical isolates with stable metronidazole resistance have been described and we evaluated the presence of pCD-METRO in the assembled genome sequences from these strains using BLAST.^23, 37^ We failed to identify pCD-METRO in the draft genome of a toxigenic NAP1 isolate that acquired stable metronidazole resistance through serial passaging under selection.^37^ We did identify pCD-METRO (fragmented over multiple contigs) in the draft genome a non-toxigenic Spanish RT010 strain with stable metronidazole resistance (strain 7032989), whereas neither the reduced-susceptible strain nor the susceptible strain from the same study contained the plasmid.^23^ We confirmed these results using PCR, as described for strain IB138 (figure 3A; lanes SP), demonstrating pCD-METRO is indeed present in strain 7032989. These data show that the presence of pCD-METRO may explain at least part of the cases of metronidazole resistance described in literature. We did not detect pCD-METRO in the sequence read archive in entries labelled as *C. difficile,* or otherwise.

**Figure 3.**
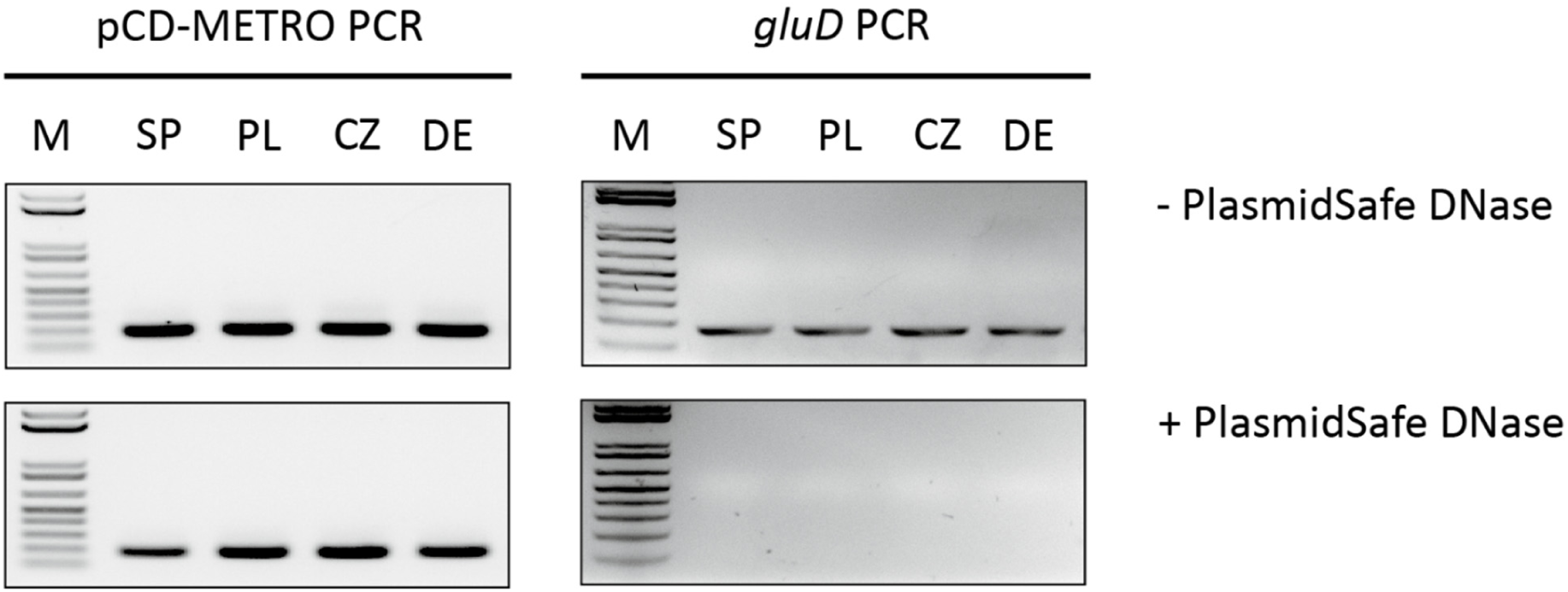
pCD-METRO is internationally disseminated. PCR analysis of strains 7032989 (RT010, Spain) (SP), 26188 (RT027, Poland) (PL), LUMCMM19 0880 (RT010, Czech Republic) (CZ) and P016134 (RT010, Germany) (DE). The product of plasmid specific amplification (targeting ORF8) or chromosomal specific amplification (*gluD*) before and after PlasmidSafe DNase treatment are shown.

Our observations above raise the question how prevalent pCD-METRO is in MTZ^R^ *C. difficile* isolates and if there is a bias towards specific types or geographic origins. As metronidazole resistance in *C. difficile* is rare, we expanded our collection of clinical isolates through our network (including the ECDC) and with selected strains from the Tolevamer and MODIFY clinical trials.^24–26^ To correct for interlaboratory differences in typing and antimicrobial susceptibility testing, all strains were retyped by capillary ribotyping and tested for metronidazole resistance using agar dilution according to CLSI guidelines in our laboratory with inclusion of appropriate control strains.^28, 29^ Although these strains, with the exception of the Tolevamer strains, were characterized as having altered metronidazole susceptibility by the senders (n=122), agar dilution performed in our own laboratory classified nearly all of these strains as metronidazole susceptible (MIC <2 mg/L). We expected pCD-METRO to be present in MTZ^R^ strains, but not in MTZ^S^ strains.

We identified three additional metronidazole resistant strains: a RT027 isolate from Poland (26188; MIC>8 mg/L), a RT010 isolate from the Czech Republic (LUMCMM19 0880; MIC>8 mg/L) and a RT010 isolate from Germany (P016134; MIC>8 mg/L) (table 1). A PCR on PSD-treated chromosomal DNA isolated from these strains yielded a positive signal using primers targeting the plasmid, but not the chromosome (figure 3), demonstrating all three strains contain pCD-METRO. We also screened our laboratory collection of RT010 strains from human and animal sources and identified 7 more MTZ^R^ strains (as determined by both agar dilution and E-test), 6 of which were positive for pCD-METRO (86%; supplemental table 1). A single RT010 strain (LUMCMM19 0830) tested MTZ^R^ resistant in agar dilution (MIC=4mg/L), but this strain was negative for pCD-METRO. Thus, all MTZ^R^ strains with an MIC≥8mg/mL identified here were found to contain pCD-METRO (22/22). By contrast, all susceptible isolates (n=562) lacked pCD-METRO.

Taken together, our results shows that pCD-METRO is internationally disseminated and can explain metronidazole resistance in both non-toxigenic- and toxigenic isolates of *C. difficile*, including those belonging to epidemic ribotypes such as RT027.

### pCD-METRO is acquired through horizontal gene transfer

Our whole genome sequence analysis revealed the acquisition of pCD-METRO by a toxigenic RT020 strain during treatment of rCDI. We made use of longitudinal fecal samples that were stored during treatment to investigate the presence of pCD-METRO in total fecal DNA at various timepoints. Total DNA derived from the fecal sample harboring the MTZ^S^ RT020 was positive for the presence of pCD-METRO (figure 4). This indicates that pCD-METRO was present in the gut reservoir of the patient. Post-FMT, pCD-METRO was no longer detected in total fecal DNA, suggesting that the fecal transplant reduced levels of pCD-METRO containing *C. difficile* and/or the donor organism to below the limit of detection of the assay. Fecal samples were stored in the absence of cryoprotectant and as a result we were unable to reculture the possible donor organism.

**Figure 4.**
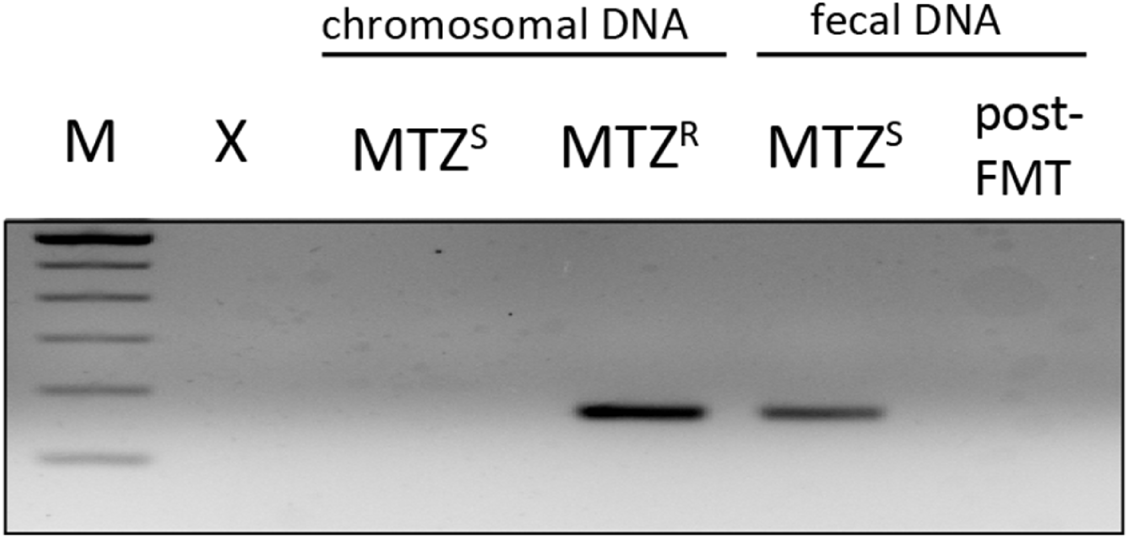
pCD-METRO is detectable in fecal total DNA. pCD-METRO is detectable in fecal total DNA from the same sample from which a MTZ^S^ RT020 *C. difficile* was isolated. Shown are the results from a PCR targeting ORF6, but similar results were obtained for other plasmid-specific primer sets (data not shown).

Though we cannot exclude the possibility that the MTZ^R^ RT020 strain was already present at the moment the MTZ^S^ RT020 strain was isolated, our results indicate that pCD-METRO was most likely acquired through horizontal gene transfer between the MTZ^S^ *C. difficile* strain and an as-of-yet uncharacterized donor organism in the gut of the patient.

### pCD-METRO confers metronidazole resistance in *C. difficile*

Above, we have clearly established a correlation between the presence of pCD-METRO and metronidazole resistance. Next, we sought to unambiguously demonstrate that acquisition of pCD-METRO, and not any secondary events, lead to metronidazole resistance. To generate isogenic strains with or without pCD-METRO, we introduced a shuttle module in the unique HaeIII restriction site of the plasmid and introduced the resulting vector, pCD-METRO^shuttle^ (pIB86; supplemental figure 1 and methods in appendix), into the RT012 laboratory strain 630Δ*erm* using standard methods.^44^ Metronidazole E-tests showed a reproducible 15-to-20-fold increase in the MIC from 0·064/0·19 mg/L for the strain without pCD-METRO^shuttle^ (figure 5) to 2-4 mg/L for the strain with pCD-METRO^shuttle^ (figure 5). These results were confirmed using agar dilution, that showed an >24-fold increase from 0·125-0.25 mg/L to 8 mg/L or higher upon introduction of pCD-METRO^shuttle^ (table 1).

**Figure 5:**
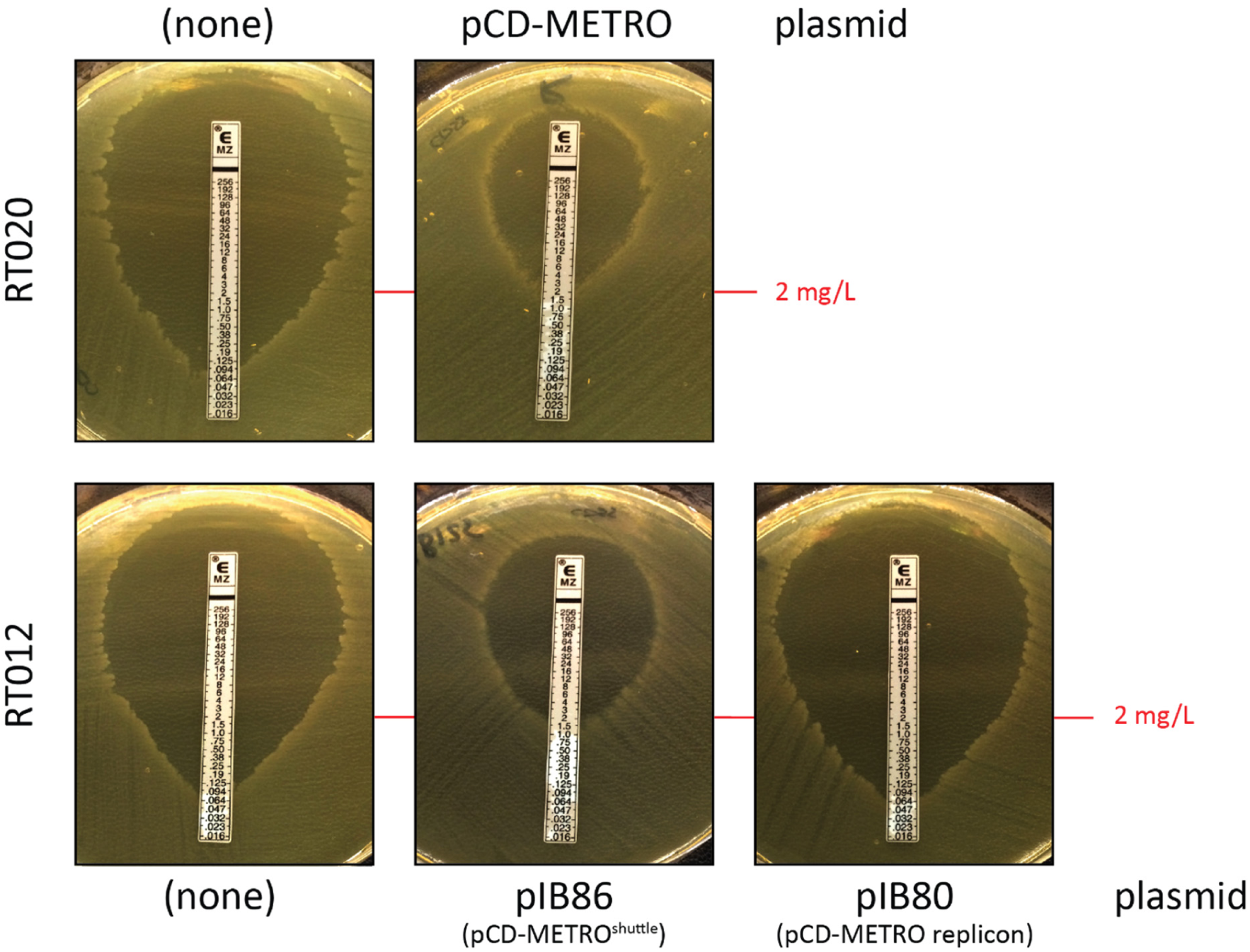
pCD-METRO confers metronidazole resistance. RT020 without plasmid (MTZ^S^, strain IB132), RT020 with pCD-METRO (MTZ^R^, strain IB133), RT012 without plasmid (MTZ^S^, strain 630Δ*erm*), RT012 with pIB86 (pCD-METRO^shuttle^, MTZ^R^, strain IB125), RT012 with pIB80 (MTZ^S^, IB90; pIB80 contains the pCD-METRO replicon but lacking the other ORFs of pCD-METRO). IB90 and IB125 are 630Δ*erm*-derivatives. E-tests were performed on BHI agar plates with CDSS. Identical results were obtained on plates without CDSS (data not shown).

As controls, we included the MTZ^S^ (IB132) and a MTZ^R^ (IB133) RT020 strain isolated from the patient. In agreement with the MIC values determined by agar dilution (MIC=0·25 mg/L and MIC=8 mg/L), these isolates showed a MIC corresponding to those observed for the MTZ^S^ and MTZ^R^ RT012 isolates, respectively (figure 5).

Overall, our results show that acquisition of pCD-METRO is sufficient to raise the MIC of *C. difficile* to values greater than the epidemiological cut-off value defined by EUCAST.^27^

### pCD-METRO contains a high copy number replicon

Read depth of pCD-METRO in our WGS data indicates an estimated copy number of 100-200, in stark contrast with the pCD6 replicon commonly used in shuttle vectors for *C. difficile* (copy number 4-10).^34^ We wanted to establish the functionality of the predicted replicon and determine the copy number sustained by this replicon in RT012 strains.

A pRPF185-based vector (pIB80) was constructed in which the conventional pCD6 replicon was replaced by a 2-kb DNA fragment of pCD-METRO that includes ORF5, encoding the putative replication protein (supplemental figure 3, appendix). Transconjugants containing this vector were readily obtained in the RT012 laboratory strain 630Δ*erm*, demonstrating this region contains a functional replicon.

Next, we compared the relative copy number of the plasmids in overnight cultures by qPCR.^34^ Based on the ratio of plasmid-locus *catP* to the chromosomal locus *rpoB,* the copy number of pCD6-replicon vector was approximately 4, concordant with results of others.^34^ By contrast, the copy number of vectors with the pCD-METRO replicon ranges from approximately 25 (for pIB80, in IB90) to 38 (pCD-METRO^shuttle^, in IB125) (figure 6). In line with these results, a strain harboring a *catP*-containing plasmid with the pCD-METRO replicon demonstrates a growth advantage over a strain harboring a similar plasmid with the pCD6 replicon when exposed to high levels (256 μg/mL) of thiamphenicol (supplemental figure 2, appendix). As pIB90 containing strains are not MTZ^R^ (figure 5), resistance to metronidazole is not mediated by a higher copy number plasmid per se, but is dependent on a determinant specific to pCD-METRO.

**Figure 6:**
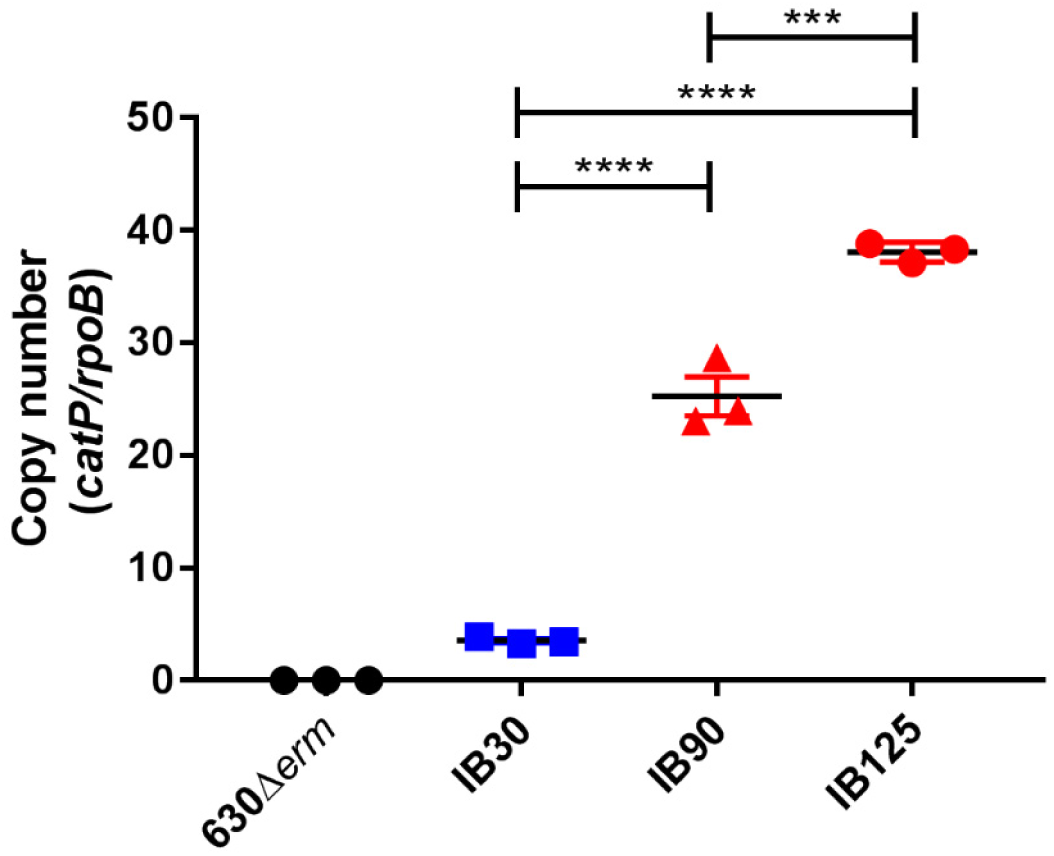
The pCD-METRO replicon sustains a high plasmid copy number. 630Δ*erm* is the wild type RT012 laboratory strain. IB30: 630Δ*erm* + pIB20 (contains pCD6 replicon); IB90: 630Δ*erm* + pIB80 (contains pCD-METRO replicon); IB125: 630Δ*erm* + pCD-METRO^shuttle^ (pIB86, contains pCD-METRO replicon). *** p<0,001, **** p<0,0001. Copy number is determined as the ratio of a plasmid locus (*catP*) relative to a chromosomal locus (*rpoB*) as determined by qPCR on total DNA. Data from strains containing a plasmid with the pCD6-replicon are indicated in blue, data from strains containing a plasmid with the pCD-METRO replicon are indicated in red. Experiments were performed in triplicate on three different technical replicates.

A difference between the read-depth estimate and the qPCR can be explained by technical bias or differences in strain background. Nevertheless, our experiments clearly demonstrate that the pCD-METRO replicon sustains plasmid levels that are approximately 10-fold greater than that of currently used replicons.

Together, these results demonstrate that pCD-METRO encodes a functional replicon that is responsible for a high copy number in *C. difficile*.

## Discussion

In this study we describe the first plasmid linked to clinically relevant antimicrobial resistance in *C. difficile*. We show that the high-copy number plasmid pCD-METRO is internationally disseminated, present in diverse PCR ribotypes - including those known to cause outbreaks – and we provide evidence for the horizontal transmission of the plasmid.

Though the presence of plasmids in *C. difficile* has been known for many years, no phenotypes associated with plasmid carriage have been described.^40, 45^ We show that introduction of pCD-METRO in susceptible strains leads to stable metronidazole resistance. Plasmids may play a broader role in antimicrobial resistance of *C. difficile*. A putative plasmid containing the aminoglycoside/linezolid resistance gene *cfrC* was recently identified *in silico*, but in contrast to our work no experiments were presented to verify the contig was in fact a plasmid conferring resistance.^46^ The presence of an antimicrobial resistance gene does not always result in resistance, and DNA-based identification of putative resistance genes without phenotypic confirmation may lead to an overestimation of the resistance frequencies.^14, 47, 48^

At present, it is unknown which gene(s) on pCD-METRO are responsible for metronidazole resistance. Nitroimidazole reductase (*nim*) genes have been implicated in resistance to nitroimidazole type antibiotics.^22^ Though the presence of a truncated *nim* gene on pCD-METRO is intriguing, we do not believe this gene to be responsible for the phenotype for several reasons. Structural modelling of the predicted protein shows that it lacks the catalytic domain, and introduction of the ORF under the control of an inducible promoter did not confer resistance in our laboratory strain (data not shown). Moreover, the RT027 strain R20291 encodes a putative 5-nitroimidazole reductase (R20291_1308) and is not resistant to metronidazole, implying the presence of a *nim* gene is not causally related to metronidazole resistance in *C. difficile*. Further research is necessary to determine the mechanism for metronidazole resistance in *C. difficile* conferred by pCD-METRO, and to investigate the contribution of the high copy number (figure 6) to the resistance phenotype.

Our work, combined with that of others, suggests that metronidazole resistance is multifactorial and other factors than pCD-METRO can cause or contribute to metronidazole resistance in *C. difficile*. For instance, pCD-METRO may not explain low level resistance, heterogeneous resistance, or stable resistance resulting from serial passaging of isolated strains under metronidazole selection.^23, 37, 49^ We also observed that MIC values in agar dilution experiments differed between MTZ^R^ isolates of different PCR ribotypes despite carriage of pCD-METRO, suggesting a contribution of chromosomal or other extrachromosomal loci to absolute resistance levels. Though the SNP we identified in the MTZ^R^ RT020 strain was not found in the MTZ^R^ RT010, we cannot exclude that it contributes to the resistance in the patient strain. Notably, all natural isolates of *C. difficile* with a MIC ≥8mg/L tested positive for pCD-METRO, whereas a plasmid-negative MTZ^R^ strain showed MICs below these levels (MIC=4 mg/L).

The pCD-METRO plasmid appears to be internationally disseminated (table 1), although further research is necessary to determine how prevalent the plasmid is in metronidazole resistant *C. difficile* isolates. This study attempted to enrich for metronidazole resistant strains as this resistance is scarce in *C. difficile*. We received strains which were reported to be metronidazole resistant by the senders. However, when performing antimicrobial susceptibility testing for these strains with agar dilution in our own laboratory, virtually all strains had MIC values below the epidemiological cut-off value from EUCAST for metronidazole and were considered susceptible. It is not entirely clear how these differences came into existence. Depending on handling of the sample material and freeze-thawing cycles, it is possible that inducible metronidazole resistance, unrelated to pCD-METRO, was initially measured and that this was lost after storage and lack of selection. For this reason we ended up having very few metronidazole resistant isolates of other PCR ribotypes than RT010 (RT020 and RT027).

The pCD-METRO plasmid appears to be transmissible (figures 1, 2 and 4). Horizontal gene transfer is consistent with the observed high level of sequence conservation between the RT010 and RT020 pCD-METRO plasmids sequenced in this study. Nevertheless, we failed to demonstrate intraspecies transfer with different donor and recipient strains of *C. difficile* under laboratory conditions (appendix), suggesting that the strains tested (or possible the species) lack a determinant required for transfer. Together with its size and the presence of mobilization genes (figure 2A), we therefore hypothesize that pCD-METRO is mobilizable from an uncharacterized donor organism.^50^ We screened the complete sequence read archive of the NCBI (paired-end Illumina data) for potential sources of the plasmid, but failed to identify any entries with reliable mapping (>1% of data) to pCD-METRO (data not shown).

As more reports are published associating metronidazole with higher disease recurrence and treatment failure, a shift in consensus for using metronidazole as first line treatment for mild to moderate CDI is occurring.^51^ The reason for treatment failure is currently unknown, but no correlation between MTZ^R^ *C. difficile* isolates and treatment failure seems to exist.^47^ We also observed that clinical isolates from subjects in which metronidazole treatment failed, were metronidazole susceptible and pCD-METRO negative (supplemental table 1, appendix).^25^ These observations, however, do not rule out a role for (other) metronidazole resistant organisms, potentially harboring pCD-METRO, in treatment failure. Indeed, levels of metronidazole at the end of the colon and in fecal material are low, and members of the microbiota involved in inactivation or sequestering of metronidazole may allow for growth of MTZ^S^ species.^52–55^

Our observation of a transmissible plasmid associated with metronidazole resistance in *C. difficile* and the gut microbiome has implications for clinical practice. First, it warrants a further investigation into the role of the plasmid in metronidazole treatment failure in CDI. Second, though this work can be seen as one more argument against the use of metronidazole as a first line treatment of CDI, detection of the plasmid in fecal material might also guide treatment decisions. And finally, screening of donors of fecal material intended for FMT might be desirable to reduce the possibility of transferring pCD-METRO to *C. difficile* in patients.

## Acknowledgements

This work was supported, in part, by a VIDI fellowship (864.13.003) from the Netherlands Organization for Scientific Research, a Gisela Thier Fellowship from the Leiden University Medical Center, and intramural funds to WKS. We would like to thank P. Spigaglia, M. Krutova, R. Peetso, M. Patyi, E. Nováková, E. Piepenbrock, S. Johnson and MSD for strains, and the ECDC for supporting the typing of MTZ^R^ strains. We would also like to thank P. Bredenbeek for help in constructing pCD-METRO^shuttle^.

## Author statements

Performed experiments: IMB, ES, CH, IMJGBS, WKS. Analyzed data: BVHH, IMB, WKS, JC. Contributed patient samples and metadata: EMT, EJK. Contributed reagents: RB, BVHH. Drafted manuscript: IMB, BVHH, JC, EJK, WKS. All authors edited and approved the final version of the manuscript.

## Appendix

- Supplemental methods
- Supplemental results
- Supplemental figure 1: pIB86 (pCD-METRO^shuttle^) plasmid map
- Supplemental figure 2: Growth curve of strains carrying different replicons in the presence of thiamphenicol
- Supplemental figure 3: pIB90 plasmid map
- Supplemental table 1: Overview of all strains tested
- Supplemental table 2: Plasmids used in this study
- Supplemental table 3: Oligonucleotides used

