## Supplemental Information for "Plasmid-mediated metronidazole resistance in *Clostridioides difficile*"

[1] Department of Medical Microbiology, Center for Infectious Diseases, Leiden University Medical Center, Leiden, The Netherlands; [2] Centre for Microbial Cell Biology, Leiden, the Netherlands; [3] Netherlands Centre for One Health, the Netherlands; [4] Center for Microbiome Analyses and Therapeutics, Leiden University Medical Center, Leiden, the Netherlands; [5] Departamento de Patología Animal, Facultad de Veterinaria, Universidad Zaragoza, Zaragoza, Spain; [6] National Institute for Public Health and the Environment, Bilthoven, the Netherlands.

Running title: Metronidazole resistance in *C. difficile*

### Supplemental methods

#### Whole Genome Sequencing and SNP analysis

Nine mL of overnight cultures were centrifuged at 4000g for 10 minutes and the pellet was resuspended in 800 µl of PBS. 24 µl lysozyme (Thermo Fisher Scientific) with a concentration of 50 g/L was added and incubated for 20 minutes at 37°C. Afterwards 15 µl proteinase K (Qiagen, The Netherlands) with a concentration of 20 g/L was added and incubated for 20 minutes at 37°C. DNA was extracted on a QIAsymphony (Qiagen, The Netherlands) with the QIAsymphony DSP Virus/Pathogen Midi Kit according to the manufacturer's instructions.

All 18 samples (table 1) were sequenced by GenomeScan (Leiden, Netherlands) on a HiSeq4000, with paired-end settings and a read length of 150 bp.

All *C. difficile* samples isolated from the patient were assembled using an inhouse pipeline. The first step in the pipeline is the removal of sequencing TrueSeq adapters with `Cutadapt` v1.2.1,[1] afterwards quality trimming with `prinseq-lite` v0.20.0,[2] with the parameters for left-hand, right-hand and average quality trimming of 30. Additionally, all reads which included Ns or non-IUPAC characters were discarded. After quality control the assembly is performed on six different assemblers. `Spades` v3.10.1 was used with standard parameters.[3] `IDBA_UD` v1.1.1 (custom compiled, with higher kmer range) was used with a kmer-range starting from 20 to 110 and ending with 120.[4] All kmer-ranges were used with step size 10. `Ray` version 2.3.0 and `Velvet` v1.2.10 were used with a kmer-range between 51 and 151, `Abyss` v1.5.2 was used with kmers 51 and 61.[5,6] `Edena` v3.131028 was used on the non-trimmed reads with an overlap range between 76 and 146.[7,8] Additionally one assembly per assembler was attempted with the optimal kmer as predicted by `kmergenie` v1.6741 on the interleaved fastq files.[9] Interleaving was performed via the script available at https://gist.github.com/ngcrawford/2232505. Afterwards reads were mapped back to all assemblies for quality control purposes with Bowtie2 v2.3.1, and SAM files were converted to sorted and indexed BAM files with Samtools v1.5 to obtain mapping rates to the assembly.[10,11] After this step, all contigs from assemblies with a length smaller than 304 bp were discarded, as well as contigs corresponding to the phiX phage spike in (GenBank accession number J02482.1). To remove contaminating contigs, contigs from all assemblies were compared with Blastn v2.6 against the NCBI database (download July 10, 2018, standard parameters, except evalue of 0.0001). [12,13] Taxonomy was estimated with the LCA algorithm as implemented in MEGAN, except that only Blast matches with a minimum length of 100bp, and as well only matches not deviating more than 10% in length from the longest match were considered.[14] Filtering was performed on phylum level and the dominant phylum was determined by the amount of base pairs in the assigned blast matches. All contigs assigned to another phylum were discarded. After this step, reads were mapped and assemblies were scaffolded as described previously. The insert size of the paired-end reads from the scaffolded assemblies was estimated with CollectInsertSizeMetrics from the Picard tools package v1.94 (http://broadinstitute.github.io/picard). This insert size estimation was used for gapfilling with GapFiller v1.11 and 20 iterations of gapfilling, followed again by read mapping.[15] For quality control, the expected genome size was estimated with kmerspectrumanalyzer download August 2013 and Jellyfish v1.1.11. [16,17] Using bedtools genomecov v2.2.16 the read coverage of the assemblies were calculated.[18] All sequence ranges larger than 20 bp with less than 50% coverage and with a larger distance than 200 bp from the contig end were manually inspected, unless only Ns were contained in the sequence. Final evaluation was performed with the values for N50, assembly size (in relation to predicted genome size), mapping rate and manual inspection of low coverage sites. The assembly being evaluated as being best was performed with the Edena assembler and an overlap of 126, with contamination and length filtering, without additional scaffolding and gapfilling.

Annotation was performed with an in-house pipeline consisting out of gene calling with `prodigal` v2.6.3, rRNA prediction with `Rnammer` v1.2, tRNA prediction with `Aragorn` v1.2.38 and CRISPR annotation with the `CRISPR recognition tool` v1.2. [19-22]. Genes predicted over an assembly gap and containing more than 50% N were discarded. Protein annotation was performed with `InterproScan` v5.26-65.0, `PRIAM` March 2015 (together with legacy `blast` v2.2.26) and `dbCAN` v5.0.[23-26] Additional EC numbers were derived via the GO terms derived from the `InterproScan` output.[27] If an EC number could be assigned to a protein sequence, then it was annotated with its canonical name. Otherwise, all `InterproScan` domain names were searched for terms relating to functions involved in virus replication, sporulation, ribosomal proteins, CRISPR or words containing any subunit. If this did not lead to any result, all domain names were searched for words ending in “ase”, indicating potential enzyme functions. Otherwise a random domain name was picked. All these annotation steps were done while disregarding generic or uninformative terms (e.g. containing “hypothetical”, “DUF”, “uncharacterized”). Domains containing these words were only considered after all other steps did not lead to any result. This annotation was furthermore manually reviewed and the annotations from the assembly of *C. difficile* 630 (based on `Blastn` comparison on gene level) were transferred where applicable.

SNP typing was performed after selecting the best reference assembly. The SNP typing was performed with the in-house pipeline `Basty` based on the `biopet` framework.[28] This pipeline performs mapping to the reference assembly with `bowtie2` v2.3.1 and SNP typing with `BCFtools` v1.1-134.[29] Groups were investigated for homozygous SNPs differentiating them. Heterozygous SNPs were discarded. Genomic comparisons between assemblies were performed with `Blastn` (standard parameters, except an evalue of 0.0001). All programs were executed with standard parameters unless otherwise specified.

**Detection of the plasmid in public databases**

5403 public paired-end Illumina datasets were downloaded from the NCBI (accession numbers see supplemental table 4). An approach similar to `placnetW` was used for plasmid detection.[30,31]

All samples were downloaded with `eutils` prefetch and converted with `fastq-dump` from the `SRA toolkit` v2.8.2-1. The optimal kmer was predicted by `kmergenie` v1.6741 on the interleaved fastq files and the assembly was performed with `Velvet` v1.2.10 .[6,9,13] The assembly graph from the `Velvet` output was created with the python `networkX` library.[32] To calculate size and coverage all headers in the `Velvet` assembly were parsed into the network. The component with the most bases was considered to be the genome, and average coverage was estimated by averaging the coverage of all contigs over the amount of contigs. All other network components were considered chromosomal if their coverage did not exceed 1.5 times the coverage of the chromosome. To reduce the amount of false positive identifications the coverage was adjusted for the number of basepairs as opposed to the amount of contigs. Additionally a Blast search was performed against the chromosome of *Clostridioides difficile* 630, and all components with more than 50% genomic content were also regarded as belonging to the chromosome.[33]

To identify plasmids similar to the pCD-METRO, a homology search was performed with `Blastn` (with an e-value of 0.0001).

A further search for non-assembled plasmid sequences was performed. All samples sequenced in paired-end mode on Illumina machines were downloaded from the NCBI with `eutils` prefetch, and mapped to the plasmid sequence with `bowtie2` v2.3.1 to the plasmid sequence. The option `--no-mixed` was used to supress incorrectly mapping pairs.

**Data accessibility**

All sequence data generated in this study has been uploaded to the European Nucleotide Archive under project PRJEB24167 with accession numbers ERR2232520- ERR2232537. The genome assembly for IB136, including the annotated sequence of pCD-METRO, can be found under accession number ERZ807316.

**Antimicrobial susceptibility testing and ribotyping**

All strains were tested for metronidazole resistance using agar dilution according to Clinical & Laboratory Standards Institute guidelines and as described previously.[34,35] Strains were characterized by PCR ribotyping and a multiplex PCR as described previously.[36,37] For epsilometer tests (E-test; BioMerieux), bacterial suspensions corresponding to 1·0 McFarland turbidity were applied on BHI agar supplemented with 0·5% yeast extract (Sigma-Aldrich) and *Clostridium difficile* Selective Supplement (CDSS, Oxoid). MIC values were read after 48 hours of incubation.

**Culture**

The plasmids and bacterial strains used are listed in supplementary tables 2 and 3, respectively. Escherichia coli cells were cultured in Luria-Bertani (LB) broth in an aerobic environment at 37$^{o}$C (shaking at 200 rpm). The LB broth was supplemented with 20 µg/ml chloramphenicol and 50 µg/ml kanamycin when required. Brain Heart Infusion (BHI, Oxoid) supplemented with 0,5% yeast extract (Sigma-Aldrich), *Clostridium difficile* Selective Supplement (CDSS, Oxoid) and 20 µg/ml thiamphenicol when appropriate, was used for routine culturing of *Clostridium difficile* strains. A Don Whitley VA-1000 workstation was used for anaerobic incubation of *C. difficile* in an atmosphere of 10% $CO_2$, 10% $H_2$ and 80% $N_2$. Liquid cultures were cultured in identical conditions shaking at 120 rpm.

**Plasmid construction**

Plasmid pIB80 was constructed by ATUM (Newark, CA) and contains a pCD-METRO derived fragment inserted in between the KpnI and NcoI sites of pRPF185.[38] pIB86 was constructed using Gibson assembly using HaeIII-linearized pCD-METRO and a fragment from pRPF185. This fragment was obtained by PCR, and contained the requirements for maintenance in, and transfer from, *E. coli*. Cesium chloride purified pCD-METRO (see below) was linearized using restriction endonuclease HaeIII. Primers oWKS-1663 and oWKS-1664 annealed on pRPF185 generating a PCR shuttle-fragment containing pBR322ori-*catP-oriT-traJ*. To assemble pIB86, 100 ng of insert was assembled against a fourfold molar excess of linearized pCD-METRO backbone using a homemade Gibson Assembly Master Mix (4 U/µl Taq Ligase (Westburg), 0.004 U/µl T5 exonuclease (New England Biolabs), 0.025 U/µl Phusion polymerase (Bioké), 5% polyethyleneglycol (PEG-8000), 10 mM MgCl2, 100 mM Tris-Cl pH=7.5, 10 mM dithiothreitol, 0·2 mM dATP, 0·2 mM dTTP, 0·2 mM dCTP, 0·2 mM dGTP, and 1 mM β-nicotinamide adenine dinucleotide) for 30 minutes at 50$^{o}$C and transformed into MDS42 cells. Transformants were screened by colony PCR using primers oBH-5 and oWKS-1387. The entire sequence of pIB86 was verified by Sanger sequencing using primers oBH-1, oBH-5, oBH-6, oBH-8, oBH-9, oBH-10, oBH-11, oBH-12, oIB-120-, oIB-121, and oIB-122, oWKS-1241-, oWKS-1383, oWKS-1388, oWKS-1537, oWKS-1539, oWKS-1540, oWKS-1574, oWKS-1656, oWKS1658, oWKS-1659, oWKS-1661, oWKS-1663, oWKS-1664 and oWKS-1678.

**Plasmid maintenance and conjugative transfer**

The *E. coli* strain DH5α was used for the maintenance of plasmids pIB76, pIB77, pIB78, pIB79, pIB80. Plasmid pIB86 was maintained in the *E. coli* strain MDS42 due to the lack of IS elements in this strain that could otherwise insert themselves into the maintained plasmid.[39] The *E. coli strain* CA434 was used for conjugative transfer of plasmids into all used *C. difficile* strains. [40,41]. Conjugation was performed as previously described.[40] Routine plasmid DNA extraction was performed using the Nucleospin Plasmid (Macherey-Nagel) mini prep kits per manufacturer's instructions. The DNeasy blood and tissue kit (Qiagen) was used for isolating *C. difficile* total DNA after incubating the cells in an enzymatic lysis buffer according to instructions of the manufacturer.

**Isolation of cloning-grade pCD-METRO plasmid preparation using the $CsCl_2$ plasmid purification method**

Plasmid pCD-METRO was extracted from 400 mL of culture containing the $MTZ^R$ strain IB138 using the Macherey-Nagel Nucleobond Xtra Midi kit. Using the $CsCl_2$ plasmid purification method this plasmid prep was further cleaned as summarized hereafter. pCD-METRO plasmid prep was added to TE buffer (10 mM Tris pH=8·0, 1 mM EDTA) and $CsCl_2$ was added to a density of 1g/g. Approximately 220 µg/ml ethidium bromide was added to this solution after which samples were spun down in a Beckman Coulter Optima XE-90 ultracentrifuge for 17 hours at 65.000 rpm, 20$^{o}$C. Bands were visualized with UV light and plasmid DNA was collected by withdrawing the lowest of the two resulting bands with an 18 gauge needle. To remove ethidium bromide 1x vol/vol 5M NaCl saturated N-butanol was used to remove the upper (purple) phase after centrifugation. Samples were ethanol precipitated twice prior to resuspending purified plasmid DNA in TE buffer.

**qPCR for copy number determination**

For the real time quantitative PCR (qPCR) experiments cells were collected from overnight (17h) cultures and lysed as described without lysozyme.[42] Total DNA was isolated using a phenol-chloroform extraction protocol.[43] Total DNA was diluted to a concentration of 10 ng/µl and 4 µl of the diluted DNA sample was added to 6 µl of a mixture containing SYBR Green Supermix (Bio-Rad) and gene-specific primers (0.4 µM total) for a total volume of 10 µl per well. Gene specific primers used were TEQ009 and TEQ010 (*rpoB*), and RP314 and RP315 (*catR*) as described in.[44] Experiments were performed in triplicates on three different technical replicates. To determine plasmid copy number results were normalized against the housekeeping gene *rpoB*.

**Conjugation of pCD-METRO between *C. difficile* strains**

A spontaneous rifampicin resistant 630Δ*erm* strain was generated by incubating cells on a BHI agar plate supplemented with 25 µg/ml rifampicin until colonies appeared. The resulting strain EVE17 was found to have acquired the well described R505K mutation in *rpoB* as verified by Sanger sequencing using primers oIB-103, oIB-104, oIB-105, oIB-106, oIB-107 and oIB-108.[45] Conjugative transfer was attempted between a pCD-METRO harboring RT010 strain IB138 and CD37, EVE17 or WKS1710 (630Δ*erm tcdA*::CT *tcdB*::CT; $Li^R$). [40]

Transconjugants were selected using 4 µg/ml metronidazole combined with either 20 µg/ml rifampicin (for CD37 and EVE17) or 20 µg/ml lincomycin (for WKS1710).

**Preparation of Figures**

Agarose gel and E-test images were acquired using a BioRad GelDoc XR, processed in Adobe Photoshop CC 2018 (Adobe). Plasmid maps were generated using Artemis and DNAplotter.[46,47] All figures were prepared for publication in Adobe Illustrator CC 2018 (Adobe).

### Supplemental results

#### Growth curve for determination of copy number effect on resistance

We hypothesized that a higher plasmid copy number would also lead to more copies of the resistance marker on the plasmid and thus to a possible increase of resistance to the corresponding antibiotic. To test this hypothesis, exponentially growing starter cultures of *C. difficile* AP38 (pCD6 replicon; copy number ~4) or IB90 (pCD-METRO replicon, copy number ~30) were inoculated to an $OD_{600}$ of 0·05 in BHI without or supplemented with 20 or 256 µg/ml thiamphenicol. No significant differences are observed for both strains when grown without antibiotic or in the presence of 20 µg/ml thiamphenicol (supplemental figure 2). During 8 hours of growth it is apparent that the lag phase of the *C. difficile* strain containing the pCD6-replicon based vector is larger than the strain harbouring the pCD-METRO-replicon based vector in the presence of 256 µg/ml of thiamphenicol (supplemental figure 2). Also, growth of AP38 is slower than the growth of IB90, as evident from the slope of the curve. Analyzing the slope on a linear scale between T=3 and T=6 using the conventional $\Delta y/\Delta x$ the slope for pCD-METRO at 256 µg/ml thiamphenicol is 0·251 versus 0·177 of pCD6. These data show that when faced with high concentrations of thiamphenicol, a higher plasmid copy number containing *catP* leads to better adaptation.

#### No demonstrable intraspecies transfer of pCD-METRO

We wanted to determine if pCD-METRO was transferable between *C. difficile* isolates. Because the pCD-METRO contains *mobA*/*mobC* homologs it is possible the plasmid is mobilizable and/or conjugative.[48] Conjugations between *C. difficile* donor and recipient strains were attempted. Strain IB138 was used as a donor as this strain only carries the pCD-

METRO plasmid without any other extrachromosomal elements.[49] A rifampicin resistant 630Δ*erm* strain and CD37, a strain frequently used in filter-mating experiments, were used as recipients for pCD-METRO using IB138 as a donor.[50] No pCD-METRO containing RT012 and RT09 transconjugants could be obtained when selecting for colonies using metronidazole and/or rifampicin. To exclude the possibility of the antibiotic selection interfering with the conjugation process, an Clostron mutant (630Δ*erm* *tcdA*::CT *tcdB*::CT) was also used as a recipient and transconjugants were selected for using lincomycin and/or metronidazole. Again, no RT012 strains containing pCD-METRO were obtained, but the RT010 donor strain did acquire spontaneous lincomycin resistance (data not shown). The lincomycin resistance mechanism has not been further investigated. Though we cannot exclude the possibility that intraspecies transfer is possible between strains that were not tested, or under conditions different from the ones we employed here, we have not been able to observe transfer of pCD-METRO from RT010 into our lab strains under the conditions used.

### Supplemental figures

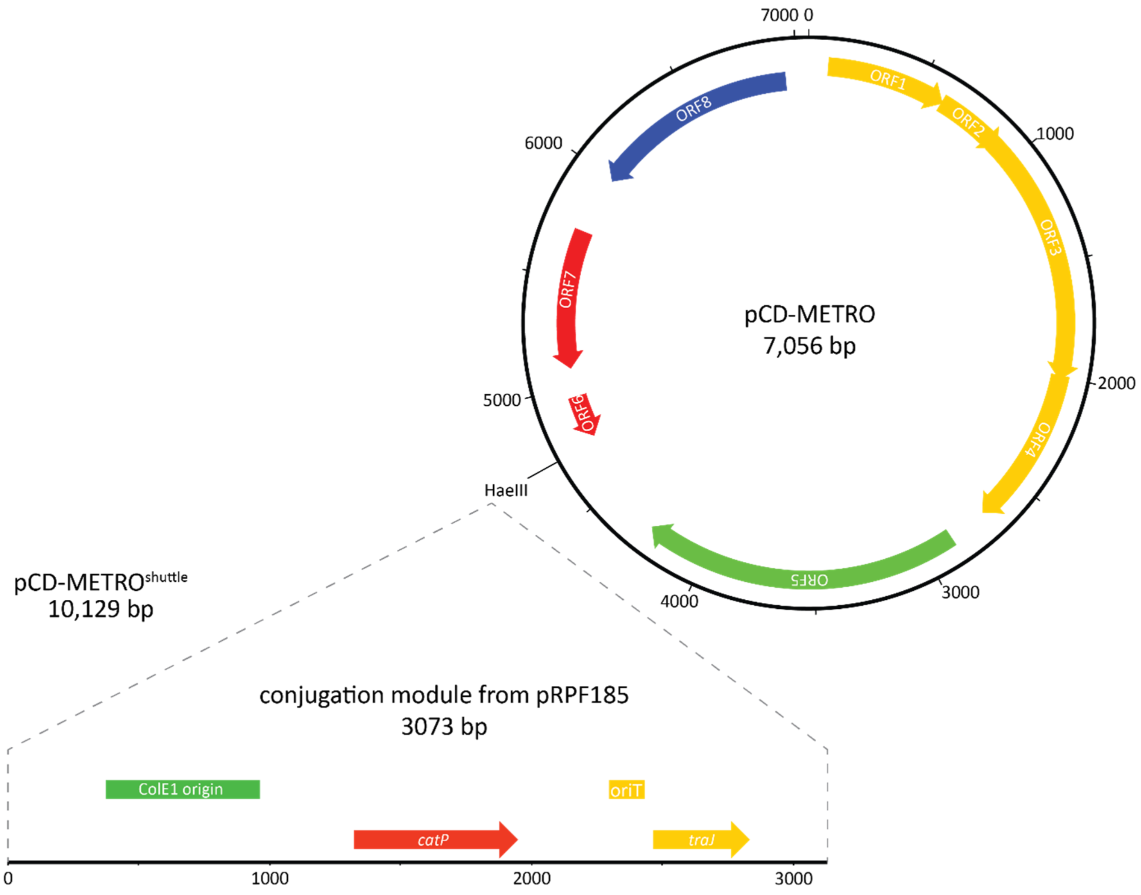


**Supplemental figure 1.** Schematic representation of the construction of pCD-METRO$^{shuttle}$ (pIB86): the PCR-amplified conjugation module from pRPF185 was inserted in the unique HaeIII restriction site through Gibson assembly.

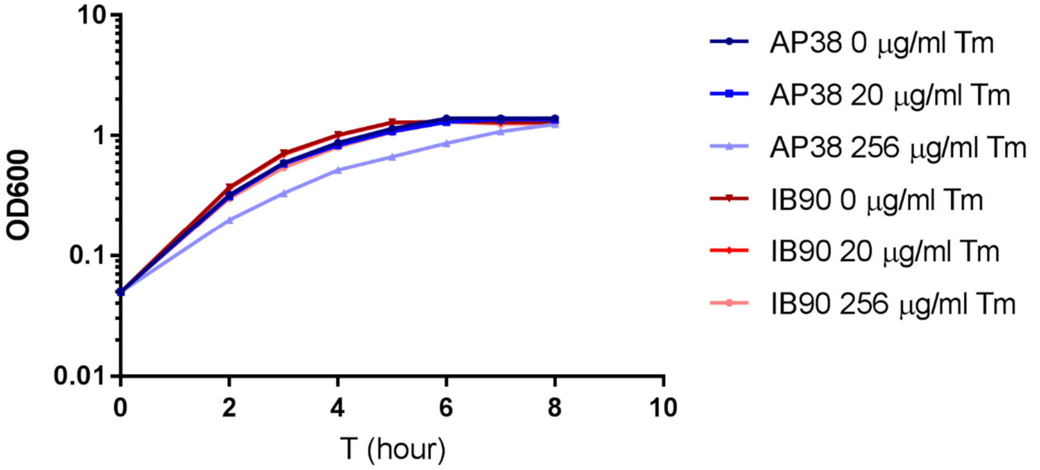


**Supplemental figure 2**. A strain derived from the laboratory RT012 strain 630Δ*erm* harboring a plasmid containing the pCD-METRO replicon (IB90, red lines) has a growth advantage over a strain containing a plasmid with the pCD6 replicon (AP38, blue lines) when cultured at high levels of thiamphenicol (Tm). AP38: 630Δ*erm* + pAP24 (pCD6 replicon); IB90: 630Δ*erm* + pIB80 (pCD-METRO replicon).

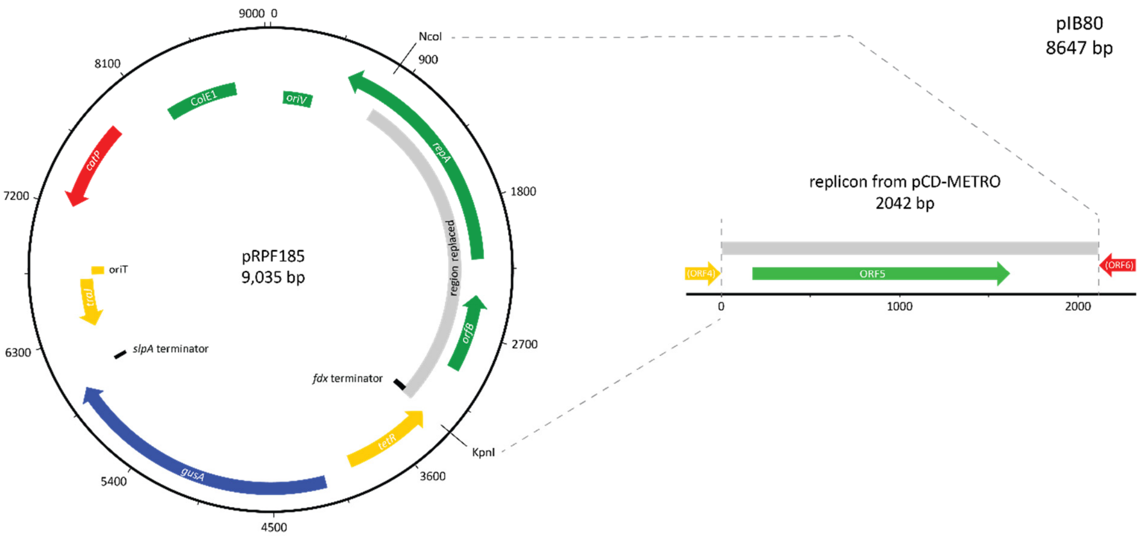


**Supplemental figure 3**: pIB80 was generated by cloning the area containing the ORF encoding the putative replication protein from pCD-METRO into the indicated area of pRPF185, disrupting the pCD6 replicon (*orfB* + *repA*).

**Strains charachterized with agar dilution and tested for presence of pCD-METRO**

| Name | Characteristics | PCR ribotype | MTZ resistance | Source | Reference |
|---|---|---|---|---|---|
| LUMCMM19 0405 | pCD-METRO - | 078 | S | Universidad de Zaragoza | This study |
| LUMCMM19 0406 | pCD-METRO - | 078 | S | Universidad de Zaragoza | This study |
| LUMCMM19 0407 | pCD-METRO - | 078 | S | Universidad de Zaragoza | This study |
| LUMCMM19 0408 | pCD-METRO - | 078 | S | Universidad de Zaragoza | This study |
| LUMCMM19 0409 | pCD-METRO - | 078 | S | Universidad de Zaragoza | This study |
| LUMCMM19 0410 | pCD-METRO - | 078 | S | Universidad de Zaragoza | This study |
| LUMCMM19 0411 | pCD-METRO - | 078 | S | Universidad de Zaragoza | This study |
| LUMCMM19 0412 | pCD-METRO - | 078 | S | Universidad de Zaragoza | This study |
| LUMCMM19 0413 | pCD-METRO - | 078 | S | Universidad de Zaragoza | This study |
| LUMCMM19 0414 | pCD-METRO - | 078 | S | Universidad de Zaragoza | This study |
| LUMCMM19 0415 | pCD-METRO - | 078 | S | Universidad de Zaragoza | This study |
| LUMCMM19 0416 | pCD-METRO - | 078 | S | Universidad de Zaragoza | This study |
| LUMCMM19 0417 | pCD-METRO - | 078 | S | Universidad de Zaragoza | This study |
| LUMCMM19 0418 | pCD-METRO - | 078 | S | Universidad de Zaragoza | This study |
| LUMCMM19 0419 | pCD-METRO - | 078 | S | Universidad de Zaragoza | This study |
| LUMCMM19 0420 | pCD-METRO - | 078 | S | Universidad de Zaragoza | This study |
| LUMCMM19 0421 | pCD-METRO - | 078 | S | Universidad de Zaragoza | This study |
| LUMCMM19 0422 | pCD-METRO - | 078 | S | Universidad de Zaragoza | This study |
| LUMCMM19 0423 | pCD-METRO - | 078 | S | Universidad de Zaragoza | This study |
| LUMCMM19 0424 | pCD-METRO - | 078 | S | Universidad de Zaragoza | This study |
| LUMCMM19 0425 | pCD-METRO - | 078 | S | Universidad de Zaragoza | This study |
| LUMCMM19 0426 | pCD-METRO - | 078 | S | Universidad de Zaragoza | This study |
| LUMCMM19 0427 | pCD-METRO - | 078 | S | Universidad de Zaragoza | This study |
| LUMCMM19 0428 | pCD-METRO - | 078 | S | Universidad de Zaragoza | This study |
| LUMCMM19 0429 | pCD-METRO - | 078 | S | Universidad de Zaragoza | This study |
| LUMCMM19 0430 | pCD-METRO - | 078 | S | Universidad de Zaragoza | This study |
| LUMCMM19 0431 | pCD-METRO - | 078 | S | Universidad de Zaragoza | This study |
| LUMCMM19 0432 | pCD-METRO - | 078 | S | Universidad de Zaragoza | This study |
| LUMCMM19 0433 | pCD-METRO - | 078 | S | Universidad de Zaragoza | This study |
| LUMCMM19 0434 | pCD-METRO - | 078 | S | Universidad de Zaragoza | This study |
| LUMCMM19 0435 | pCD-METRO - | 078 | S | Universidad de Zaragoza | This study |
| LUMCMM19 0436 | pCD-METRO - | 078 | S | Universidad de Zaragoza | This study |
| LUMCMM19 0437 | pCD-METRO - | 078 | S | Universidad de Zaragoza | This study |
| LUMCMM19 0438 | pCD-METRO - | 078 | S | Universidad de Zaragoza | This study |
| LUMCMM19 0439 | pCD-METRO - | 078 | S | Universidad de Zaragoza | This study |
| LUMCMM19 0440 | pCD-METRO - | 078 | S | Universidad de Zaragoza | This study |
| LUMCMM19 0441 | pCD-METRO - | 078 | S | Universidad de Zaragoza | This study |
| LUMCMM19 0442 | pCD-METRO - | 078 | S | Universidad de Zaragoza | This study |
| LUMCMM19 0443 | pCD-METRO - | 078 | S | Universidad de Zaragoza | This study |
| LUMCMM19 0444 | pCD-METRO - | 078 | S | Universidad de Zaragoza | This study |
| LUMCMM19 0445 | pCD-METRO - | 078 | S | Universidad de Zaragoza | This study |
| LUMCMM19 0446 | pCD-METRO - | 078 | S | Universidad de Zaragoza | This study |
| LUMCMM19 0447 | pCD-METRO - | 078 | S | Universidad de Zaragoza | This study |
| LUMCMM19 0448 | pCD-METRO - | 078 | S | Universidad de Zaragoza | This study |
| LUMCMM19 0449 | pCD-METRO - | 078 | S | Universidad de Zaragoza | This study |
| LUMCMM19 0450 | pCD-METRO - | 078 | S | Universidad de Zaragoza | This study |
| LUMCMM19 0451 | pCD-METRO - | 078 | S | Universidad de Zaragoza | This study |
| LUMCMM19 0452 | pCD-METRO - | 078 | S | Universidad de Zaragoza | This study |
| LUMCMM19 0453 | pCD-METRO - | 078 | S | Universidad de Zaragoza | This study |
| LUMCMM19 0454 | pCD-METRO - | 078 | S | Universidad de Zaragoza | This study |
| LUMCMM19 0455 | pCD-METRO - | 078 | S | Universidad de Zaragoza | This study |
| LUMCMM19 0456 | pCD-METRO - | 078 | S | Universidad de Zaragoza | This study |
| LUMCMM19 0457 | pCD-METRO - | 078 | S | Universidad de Zaragoza | This study |
| LUMCMM19 0458 | pCD-METRO - | 078 | S | Universidad de Zaragoza | This study |
| LUMCMM19 0459 | pCD-METRO - | 078 | S | Universidad de Zaragoza | This study |
| LUMCMM19 0460 | pCD-METRO - | 078 | S | Universidad de Zaragoza | This study |
| LUMCMM19 0461 | pCD-METRO - | 078 | S | Universidad de Zaragoza | This study |
| LUMCMM19 0462 | pCD-METRO - | 078 | S | Universidad de Zaragoza | This study |
| LUMCMM19 0463 | pCD-METRO - | 078 | S | Universidad de Zaragoza | This study |
| LUMCMM19 0464 | pCD-METRO - | 078 | S | Universidad de Zaragoza | This study |
| LUMCMM19 0465 | pCD-METRO - | 078 | S | Universidad de Zaragoza | This study |
| LUMCMM19 0466 | pCD-METRO - | 078 | S | Universidad de Zaragoza | This study |
| LUMCMM19 0467 | pCD-METRO - | 078 | S | Universidad de Zaragoza | This study |
| LUMCMM19 0468 | pCD-METRO - | 078 | S | Universidad de Zaragoza | This study |
| LUMCMM19 0469 | pCD-METRO - | 078 | S | Universidad de Zaragoza | This study |
| LUMCMM19 0470 | pCD-METRO - | 078 | S | Universidad de Zaragoza | This study |
| LUMCMM19 0471 | pCD-METRO - | 078 | S | Universidad de Zaragoza | This study |
| LUMCMM19 0472 | pCD-METRO - | 078 | S | Universidad de Zaragoza | This study |
| LUMCMM19 0473 | pCD-METRO - | 078 | S | Universidad de Zaragoza | This study |
| LUMCMM19 0474 | pCD-METRO - | 078 | S | Universidad de Zaragoza | This study |
| LUMCMM19 0475 | pCD-METRO - | 078 | S | Universidad de Zaragoza | This study |
| LUMCMM19 0476 | pCD-METRO - | 078 | S | Universidad de Zaragoza | This study |
| LUMCMM19 0477 | pCD-METRO - | 078 | S | Universidad de Zaragoza | This study |
| LUMCMM19 0478 | pCD-METRO - | 078 | S | Universidad de Zaragoza | This study |
| LUMCMM19 0479 | pCD-METRO - | 078 | S | Universidad de Zaragoza | This study |
| LUMCMM19 0480 | pCD-METRO - | 078 | S | Universidad de Zaragoza | This study |
| LUMCMM19 0481 | pCD-METRO - | 078 | S | Universidad de Zaragoza | This study |
| LUMCMM19 0482 | pCD-METRO - | 078 | S | Universidad de Zaragoza | This study |
| LUMCMM19 0483 | pCD-METRO - | 078 | S | Universidad de Zaragoza | This study |
| LUMCMM19 0484 | pCD-METRO - | 078 | S | Universidad de Zaragoza | This study |
| LUMCMM19 0485 | pCD-METRO - | 078 | S | Universidad de Zaragoza | This study |
| LUMCMM19 0486 | pCD-METRO - | 137 | S | ECDIS 2013 | MP Bauer et al., 2011 |
| LUMCMM19 0487 | pCD-METRO - | 207 | S | ECDIS 2013 | MP Bauer et al., 2011 |
| LUMCMM19 0488 | pCD-METRO - | 018 | S | ECDIS 2013 | MP Bauer et al., 2011 |
| LUMCMM19 0489 | pCD-METRO - | 014 | S | ECDIS 2013 | MP Bauer et al., 2011 |
| LUMCMM19 0490 | pCD-METRO - | 014 | S | ECDIS 2013 | MP Bauer et al., 2011 |
| LUMCMM19 0491 | pCD-METRO - | 002 | S | ECDIS 2013 | MP Bauer et al., 2011 |
| LUMCMM19 0492 | pCD-METRO - | 015 | S | ECDIS 2013 | MP Bauer et al., 2011 |
| LUMCMM19 0493 | pCD-METRO - | 014 | S | ECDIS 2013 | MP Bauer et al., 2011 |
| LUMCMM19 0494 | pCD-METRO - | 020 | S | ECDIS 2013 | MP Bauer et al., 2011 |
| LUMCMM19 0495 | pCD-METRO - | 001 | S | ECDIS 2013 | MP Bauer et al., 2011 |
| LUMCMM19 0496 | pCD-METRO - | 020 | S | ECDIS 2013 | MP Bauer et al., 2011 |
| LUMCMM19 0497 | pCD-METRO - | 258 | S | ECDIS 2013 | MP Bauer et al., 2011 |
| LUMCMM19 0498 | pCD-METRO - | 258 | S | ECDIS 2013 | MP Bauer et al., 2011 |
| LUMCMM19 0499 | pCD-METRO - | 087 | S | ECDIS 2013 | MP Bauer et al., 2011 |
| LUMCMM19 0500 | pCD-METRO - | 012 | S | ECDIS 2013 | MP Bauer et al., 2011 |
| LUMCMM19 0501 | pCD-METRO - | 014 | S | ECDIS 2013 | MP Bauer et al., 2011 |
| LUMCMM19 0502 | pCD-METRO - | 014 | S | ECDIS 2013 | MP Bauer et al., 2011 |
| LUMCMM19 0503 | pCD-METRO - | 078 | S | ECDIS 2013 | MP Bauer et al., 2011 |
| LUMCMM19 0504 | pCD-METRO - | 023 | S | ECDIS 2013 | MP Bauer et al., 2011 |
| LUMCMM19 0505 | pCD-METRO - | 078 | S | ECDIS 2013 | MP Bauer et al., 2011 |
| LUMCMM19 0506 | pCD-METRO - | 027 | S | ECDIS 2013 | MP Bauer et al., 2011 |
| LUMCMM19 0507 | pCD-METRO - | 476 | S | ECDIS 2013 | MP Bauer et al., 2011 |
| LUMCMM19 0508 | pCD-METRO - | 014 | S | ECDIS 2013 | MP Bauer et al., 2011 |
| LUMCMM19 0509 | pCD-METRO - | 014 | S | ECDIS 2013 | MP Bauer et al., 2011 |
| LUMCMM19 0510 | pCD-METRO - | 014 | S | ECDIS 2013 | MP Bauer et al., 2011 |
| LUMCMM19 0511 | pCD-METRO - | 087 | S | ECDIS 2013 | MP Bauer et al., 2011 |
| LUMCMM19 0512 | pCD-METRO - | 001 | S | ECDIS 2013 | MP Bauer et al., 2011 |
| LUMCMM19 0513 | pCD-METRO - | 011 | S | ECDIS 2013 | MP Bauer et al., 2011 |
| LUMCMM19 0514 | pCD-METRO - | 005 | S | ECDIS 2013 | MP Bauer et al., 2011 |
| LUMCMM19 0515 | pCD-METRO - | 207 | S | ECDIS 2013 | MP Bauer et al., 2011 |
| LUMCMM19 0516 | pCD-METRO - | 070 | S | ECDIS 2013 | MP Bauer et al., 2011 |
| LUMCMM19 0517 | pCD-METRO - | 020 | S | ECDIS 2013 | MP Bauer et al., 2011 |
| LUMCMM19 0518 | pCD-METRO - | 020 | S | ECDIS 2013 | MP Bauer et al., 2011 |

| | | | | | |
|---|---|---|---|---|---|
| LUMCMM19 0519 | pCD-METRO - | 078 | S | ECDIS 2013 | MP Bauer et al., 2011 |
| LUMCMM19 0520 | pCD-METRO - | 005 | S | ECDIS 2013 | MP Bauer et al., 2011 |
| LUMCMM19 0521 | pCD-METRO - | 020 | S | ECDIS 2013 | MP Bauer et al., 2011 |
| LUMCMM19 0522 | pCD-METRO - | 002 | S | ECDIS 2013 | MP Bauer et al., 2011 |
| LUMCMM19 0523 | pCD-METRO - | 087 | S | ECDIS 2013 | MP Bauer et al., 2011 |
| LUMCMM19 0524 | pCD-METRO - | NT | S | ECDIS 2013 | MP Bauer et al., 2011 |
| LUMCMM19 0525 | pCD-METRO - | 078 | S | ECDIS 2013 | MP Bauer et al., 2011 |
| LUMCMM19 0526 | pCD-METRO - | 081 | S | ECDIS 2013 | MP Bauer et al., 2011 |
| LUMCMM19 0527 | pCD-METRO - | 015 | S | ECDIS 2013 | MP Bauer et al., 2011 |
| LUMCMM19 0528 | pCD-METRO - | 045 | S | ECDIS 2013 | MP Bauer et al., 2011 |
| LUMCMM19 0529 | pCD-METRO - | 087 | S | ECDIS 2013 | MP Bauer et al., 2011 |
| LUMCMM19 0530 | pCD-METRO - | NT | S | ECDIS 2013 | MP Bauer et al., 2011 |
| LUMCMM19 0531 | pCD-METRO - | 005 | S | ECDIS 2013 | MP Bauer et al., 2011 |
| LUMCMM19 0532 | pCD-METRO - | 087 | S | ECDIS 2013 | MP Bauer et al., 2011 |
| LUMCMM19 0533 | pCD-METRO - | 027 | S | ECDIS 2013 | MP Bauer et al., 2011 |
| LUMCMM19 0534 | pCD-METRO - | 027 | S | ECDIS 2013 | MP Bauer et al., 2011 |
| LUMCMM19 0535 | pCD-METRO - | 027 | S | ECDIS 2013 | MP Bauer et al., 2011 |
| LUMCMM19 0536 | pCD-METRO - | 027 | S | ECDIS 2013 | MP Bauer et al., 2011 |
| LUMCMM19 0537 | pCD-METRO - | 027 | S | ECDIS 2013 | MP Bauer et al., 2011 |
| LUMCMM19 0538 | pCD-METRO - | 002 | S | ECDIS 2013 | MP Bauer et al., 2011 |
| LUMCMM19 0539 | pCD-METRO - | 027 | S | ECDIS 2013 | MP Bauer et al., 2011 |
| LUMCMM19 0540 | pCD-METRO - | 159 | S | ECDIS 2013 | MP Bauer et al., 2011 |
| LUMCMM19 0541 | pCD-METRO - | 002 | S | ECDIS 2013 | MP Bauer et al., 2011 |
| LUMCMM19 0542 | pCD-METRO - | 027 | S | ECDIS 2013 | MP Bauer et al., 2011 |
| LUMCMM19 0543 | pCD-METRO - | 027 | S | ECDIS 2013 | MP Bauer et al., 2011 |
| LUMCMM19 0544 | pCD-METRO - | 027 | S | ECDIS 2013 | MP Bauer et al., 2011 |
| LUMCMM19 0545 | pCD-METRO - | 027 | S | ECDIS 2013 | MP Bauer et al., 2011 |
| LUMCMM19 0546 | pCD-METRO - | 027 | S | ECDIS 2013 | MP Bauer et al., 2011 |
| LUMCMM19 0547 | pCD-METRO - | 027 | S | ECDIS 2013 | MP Bauer et al., 2011 |
| LUMCMM19 0548 | pCD-METRO - | 027 | S | ECDIS 2013 | MP Bauer et al., 2011 |
| LUMCMM19 0549 | pCD-METRO - | 027 | S | ECDIS 2013 | MP Bauer et al., 2011 |
| LUMCMM19 0550 | pCD-METRO - | 027 | S | ECDIS 2013 | MP Bauer et al., 2011 |
| LUMCMM19 0551 | pCD-METRO - | 027 | S | ECDIS 2013 | MP Bauer et al., 2011 |
| LUMCMM19 0552 | pCD-METRO - | 014 | S | ECDIS 2013 | MP Bauer et al., 2011 |
| LUMCMM19 0553 | pCD-METRO - | 029 | S | ECDIS 2013 | MP Bauer et al., 2011 |
| LUMCMM19 0554 | pCD-METRO - | 003 | S | ECDIS 2013 | MP Bauer et al., 2011 |
| LUMCMM19 0555 | pCD-METRO - | 017 | S | ECDIS 2013 | MP Bauer et al., 2011 |
| LUMCMM19 0556 | pCD-METRO - | 070 | S | ECDIS 2013 | MP Bauer et al., 2011 |
| LUMCMM19 0557 | pCD-METRO - | 023 | S | ECDIS 2013 | MP Bauer et al., 2011 |
| LUMCMM19 0558 | pCD-METRO - | 081 | S | ECDIS 2013 | MP Bauer et al., 2011 |
| LUMCMM19 0559 | pCD-METRO - | 087 | S | ECDIS 2013 | MP Bauer et al., 2011 |
| LUMCMM19 0560 | pCD-METRO - | 015 | S | ECDIS 2013 | MP Bauer et al., 2011 |
| LUMCMM19 0561 | pCD-METRO - | 014 | S | ECDIS 2013 | MP Bauer et al., 2011 |
| LUMCMM19 0562 | pCD-METRO - | 005 | S | ECDIS 2013 | MP Bauer et al., 2011 |
| LUMCMM19 0563 | pCD-METRO - | 014 | S | ECDIS 2013 | MP Bauer et al., 2011 |
| LUMCMM19 0564 | pCD-METRO - | 001 | S | ECDIS 2013 | MP Bauer et al., 2011 |
| LUMCMM19 0565 | pCD-METRO - | 070 | S | ECDIS 2013 | MP Bauer et al., 2011 |
| LUMCMM19 0566 | pCD-METRO - | 001 | S | ECDIS 2013 | MP Bauer et al., 2011 |
| LUMCMM19 0567 | pCD-METRO - | 070 | S | ECDIS 2013 | MP Bauer et al., 2011 |
| LUMCMM19 0568 | pCD-METRO - | 002 | S | ECDIS 2013 | MP Bauer et al., 2011 |
| LUMCMM19 0569 | pCD-METRO - | 014 | S | ECDIS 2013 | MP Bauer et al., 2011 |
| LUMCMM19 0570 | pCD-METRO - | 070 | S | ECDIS 2013 | MP Bauer et al., 2011 |
| LUMCMM19 0571 | pCD-METRO - | 081 | S | ECDIS 2013 | MP Bauer et al., 2011 |
| LUMCMM19 0572 | pCD-METRO - | 014 | S | ECDIS 2013 | MP Bauer et al., 2011 |
| LUMCMM19 0573 | pCD-METRO - | 015 | S | ECDIS 2013 | MP Bauer et al., 2011 |
| LUMCMM19 0574 | pCD-METRO - | 081 | S | ECDIS 2013 | MP Bauer et al., 2011 |
| LUMCMM19 0575 | pCD-METRO - | 076 | S | ECDIS 2013 | MP Bauer et al., 2011 |
| LUMCMM19 0576 | pCD-METRO - | 014 | S | ECDIS 2013 | MP Bauer et al., 2011 |
| LUMCMM19 0577 | pCD-METRO - | 017 | S | ECDIS 2013 | MP Bauer et al., 2011 |
| LUMCMM19 0578 | pCD-METRO - | 081 | S | ECDIS 2013 | MP Bauer et al., 2011 |
| LUMCMM19 0579 | pCD-METRO - | 207 | S | ECDIS 2013 | MP Bauer et al., 2011 |
| LUMCMM19 0580 | pCD-METRO - | 014 | S | ECDIS 2013 | MP Bauer et al., 2011 |
| LUMCMM19 0581 | pCD-METRO - | 050 | S | ECDIS 2013 | MP Bauer et al., 2011 |
| LUMCMM19 0582 | pCD-METRO - | 014 | S | ECDIS 2013 | MP Bauer et al., 2011 |
| LUMCMM19 0583 | pCD-METRO - | 020 | S | ECDIS 2013 | MP Bauer et al., 2011 |
| LUMCMM19 0584 | pCD-METRO - | 046 | S | ECDIS 2013 | MP Bauer et al., 2011 |
| LUMCMM19 0585 | pCD-METRO - | 012 | S | ECDIS 2013 | MP Bauer et al., 2011 |
| LUMCMM19 0586 | pCD-METRO - | 023 | S | ECDIS 2013 | MP Bauer et al., 2011 |
| LUMCMM19 0587 | pCD-METRO - | 001 | S | ECDIS 2013 | MP Bauer et al., 2011 |
| LUMCMM19 0588 | pCD-METRO - | 014 | S | ECDIS 2013 | MP Bauer et al., 2011 |
| LUMCMM19 0589 | pCD-METRO - | 001 | S | ECDIS 2013 | MP Bauer et al., 2011 |
| LUMCMM19 0590 | pCD-METRO - | 012 | S | ECDIS 2013 | MP Bauer et al., 2011 |
| LUMCMM19 0591 | pCD-METRO - | 027 | S | ECDIS 2013 | MP Bauer et al., 2011 |
| LUMCMM19 0592 | pCD-METRO - | 066 | S | ECDIS 2013 | MP Bauer et al., 2011 |
| LUMCMM19 0593 | pCD-METRO - | 017 | S | ECDIS 2013 | MP Bauer et al., 2011 |
| LUMCMM19 0594 | pCD-METRO - | 054 | S | ECDIS 2013 | MP Bauer et al., 2011 |
| LUMCMM19 0595 | pCD-METRO - | 020 | S | ECDIS 2013 | MP Bauer et al., 2011 |
| LUMCMM19 0596 | pCD-METRO - | 078 | S | ECDIS 2013 | MP Bauer et al., 2011 |
| LUMCMM19 0597 | pCD-METRO - | 012 | S | ECDIS 2013 | MP Bauer et al., 2011 |
| LUMCMM19 0598 | pCD-METRO - | 042 | S | ECDIS 2013 | MP Bauer et al., 2011 |
| LUMCMM19 0599 | pCD-METRO - | 001 | S | ECDIS 2013 | MP Bauer et al., 2011 |
| LUMCMM19 0600 | pCD-METRO - | 027 | S | ECDIS 2013 | MP Bauer et al., 2011 |
| LUMCMM19 0601 | pCD-METRO - | 027 | S | ECDIS 2013 | MP Bauer et al., 2011 |
| LUMCMM19 0602 | pCD-METRO - | 001 | S | ECDIS 2013 | MP Bauer et al., 2011 |
| LUMCMM19 0603 | pCD-METRO - | 027 | S | ECDIS 2013 | MP Bauer et al., 2011 |
| LUMCMM19 0604 | pCD-METRO - | 014 | S | ECDIS 2013 | MP Bauer et al., 2011 |
| LUMCMM19 0605 | pCD-METRO - | 001 | S | ECDIS 2013 | MP Bauer et al., 2011 |
| LUMCMM19 0606 | pCD-METRO - | 027 | S | ECDIS 2013 | MP Bauer et al., 2011 |
| LUMCMM19 0607 | pCD-METRO - | 027 | S | ECDIS 2013 | MP Bauer et al., 2011 |
| LUMCMM19 0608 | pCD-METRO - | 027 | S | ECDIS 2013 | MP Bauer et al., 2011 |
| LUMCMM19 0609 | pCD-METRO - | 027 | S | ECDIS 2013 | MP Bauer et al., 2011 |
| LUMCMM19 0610 | pCD-METRO - | 070 | S | ECDIS 2013 | MP Bauer et al., 2011 |
| LUMCMM19 0611 | pCD-METRO - | 046 | S | ECDIS 2013 | MP Bauer et al., 2011 |
| LUMCMM19 0612 | pCD-METRO - | 027 | S | ECDIS 2013 | MP Bauer et al., 2011 |
| LUMCMM19 0613 | pCD-METRO - | 135 | S | ECDIS 2013 | MP Bauer et al., 2011 |
| LUMCMM19 0614 | pCD-METRO - | 027 | S | ECDIS 2013 | MP Bauer et al., 2011 |
| LUMCMM19 0615 | pCD-METRO - | 027 | S | ECDIS 2013 | MP Bauer et al., 2011 |
| LUMCMM19 0616 | pCD-METRO - | 027 | S | ECDIS 2013 | MP Bauer et al., 2011 |
| LUMCMM19 0617 | pCD-METRO - | 027 | S | ECDIS 2013 | MP Bauer et al., 2011 |
| LUMCMM19 0618 | pCD-METRO - | 027 | S | ECDIS 2013 | MP Bauer et al., 2011 |
| LUMCMM19 0619 | pCD-METRO - | 027 | S | ECDIS 2013 | MP Bauer et al., 2011 |
| LUMCMM19 0620 | pCD-METRO - | 027 | S | ECDIS 2013 | MP Bauer et al., 2011 |
| LUMCMM19 0621 | pCD-METRO - | 001 | S | ECDIS 2013 | MP Bauer et al., 2011 |
| LUMCMM19 0622 | pCD-METRO - | 027 | S | ECDIS 2013 | MP Bauer et al., 2011 |
| LUMCMM19 0623 | pCD-METRO - | 027 | S | ECDIS 2013 | MP Bauer et al., 2011 |
| LUMCMM19 0624 | pCD-METRO - | 027 | S | ECDIS 2013 | MP Bauer et al., 2011 |
| LUMCMM19 0625 | pCD-METRO - | 027 | S | ECDIS 2013 | MP Bauer et al., 2011 |
| LUMCMM19 0626 | pCD-METRO - | 027 | S | ECDIS 2013 | MP Bauer et al., 2011 |
| LUMCMM19 0627 | pCD-METRO - | 027 | S | ECDIS 2013 | MP Bauer et al., 2011 |
| LUMCMM19 0628 | pCD-METRO - | 027 | S | ECDIS 2013 | MP Bauer et al., 2011 |
| LUMCMM19 0629 | pCD-METRO - | 027 | S | ECDIS 2013 | MP Bauer et al., 2011 |
| LUMCMM19 0630 | pCD-METRO - | 027 | S | ECDIS 2013 | MP Bauer et al., 2011 |
| LUMCMM19 0631 | pCD-METRO - | 027 | S | ECDIS 2013 | MP Bauer et al., 2011 |
| LUMCMM19 0632 | pCD-METRO - | 027 | S | ECDIS 2013 | MP Bauer et al., 2011 |
| LUMCMM19 0633 | pCD-METRO - | 027 | S | ECDIS 2013 | MP Bauer et al., 2011 |
| LUMCMM19 0634 | pCD-METRO - | 027 | S | ECDIS 2013 | MP Bauer et al., 2011 |

| | | | | | |
|---|---|---|---|---|---|
| LUMCMM19 0635 | pCD-METRO - | 027 | S | ECDIS 2013 | MP Bauer et al., 2011 |
| LUMCMM19 0636 | pCD-METRO - | 015 | S | ECDIS 2013 | MP Bauer et al., 2011 |
| LUMCMM19 0637 | pCD-METRO - | 005 | S | ECDIS 2013 | MP Bauer et al., 2011 |
| LUMCMM19 0638 | pCD-METRO - | 027 | S | ECDIS 2013 | MP Bauer et al., 2011 |
| LUMCMM19 0639 | pCD-METRO - | 027 | S | ECDIS 2013 | MP Bauer et al., 2011 |
| LUMCMM19 0640 | pCD-METRO - | 027 | S | ECDIS 2013 | MP Bauer et al., 2011 |
| LUMCMM19 0641 | pCD-METRO - | 002 | S | ECDIS 2013 | MP Bauer et al., 2011 |
| LUMCMM19 0642 | pCD-METRO - | 015 | S | ECDIS 2013 | MP Bauer et al., 2011 |
| LUMCMM19 0643 | pCD-METRO - | 001 | S | ECDIS 2013 | MP Bauer et al., 2011 |
| LUMCMM19 0644 | pCD-METRO - | 027 | S | ECDIS 2013 | MP Bauer et al., 2011 |
| LUMCMM19 0645 | pCD-METRO - | 027 | S | ECDIS 2013 | MP Bauer et al., 2011 |
| LUMCMM19 0646 | pCD-METRO - | 027 | S | ECDIS 2013 | MP Bauer et al., 2011 |
| LUMCMM19 0647 | pCD-METRO - | 027 | S | ECDIS 2013 | MP Bauer et al., 2011 |
| LUMCMM19 0648 | pCD-METRO - | 027 | S | ECDIS 2013 | MP Bauer et al., 2011 |
| LUMCMM19 0649 | pCD-METRO - | 027 | S | ECDIS 2013 | MP Bauer et al., 2011 |
| LUMCMM19 0650 | pCD-METRO - | 014 | S | ECDIS 2013 | MP Bauer et al., 2011 |
| LUMCMM19 0651 | pCD-METRO - | 027 | S | ECDIS 2013 | MP Bauer et al., 2011 |
| LUMCMM19 0652 | pCD-METRO - | 010 | S | ECDIS 2013 | MP Bauer et al., 2011 |
| LUMCMM19 0653 | pCD-METRO - | 027 | S | ECDIS 2013 | MP Bauer et al., 2011 |
| LUMCMM19 0654 | pCD-METRO - | 027 | S | ECDIS 2013 | MP Bauer et al., 2011 |
| LUMCMM19 0655 | pCD-METRO - | 027 | S | ECDIS 2013 | MP Bauer et al., 2011 |
| LUMCMM19 0656 | pCD-METRO - | 027 | S | ECDIS 2013 | MP Bauer et al., 2011 |
| LUMCMM19 0657 | pCD-METRO - | 027 | S | ECDIS 2013 | MP Bauer et al., 2011 |
| LUMCMM19 0658 | pCD-METRO - | 027 | S | ECDIS 2013 | MP Bauer et al., 2011 |
| LUMCMM19 0659 | pCD-METRO - | 027 | S | ECDIS 2013 | MP Bauer et al., 2011 |
| LUMCMM19 0660 | pCD-METRO - | 027 | S | ECDIS 2013 | MP Bauer et al., 2011 |
| LUMCMM19 0661 | pCD-METRO - | 027 | S | ECDIS 2013 | MP Bauer et al., 2011 |
| LUMCMM19 0662 | pCD-METRO - | 027 | S | ECDIS 2013 | MP Bauer et al., 2011 |
| LUMCMM19 0663 | pCD-METRO - | 027 | S | ECDIS 2013 | MP Bauer et al., 2011 |
| LUMCMM19 0664 | pCD-METRO - | 027 | S | ECDIS 2013 | MP Bauer et al., 2011 |
| LUMCMM19 0665 | pCD-METRO - | 027 | S | ECDIS 2013 | MP Bauer et al., 2011 |
| LUMCMM19 0666 | pCD-METRO - | 027 | S | ECDIS 2013 | MP Bauer et al., 2011 |
| LUMCMM19 0667 | pCD-METRO - | 027 | S | ECDIS 2013 | MP Bauer et al., 2011 |
| LUMCMM19 0668 | pCD-METRO - | 027 | S | ECDIS 2013 | MP Bauer et al., 2011 |
| LUMCMM19 0669 | pCD-METRO - | 027 | S | ECDIS 2013 | MP Bauer et al., 2011 |
| LUMCMM19 0670 | pCD-METRO - | 027 | S | ECDIS 2013 | MP Bauer et al., 2011 |
| LUMCMM19 0671 | pCD-METRO - | 027 | S | ECDIS 2013 | MP Bauer et al., 2011 |
| LUMCMM19 0672 | pCD-METRO - | 103 | S | ECDIS 2013 | MP Bauer et al., 2011 |
| LUMCMM19 0673 | pCD-METRO - | 027 | S | ECDIS 2013 | MP Bauer et al., 2011 |
| LUMCMM19 0674 | pCD-METRO - | 002 | S | ECDIS 2013 | MP Bauer et al., 2011 |
| LUMCMM19 0675 | pCD-METRO - | 045 | S | ECDIS 2013 | MP Bauer et al., 2011 |
| LUMCMM19 0676 | pCD-METRO - | 176 | S | ECDIS 2013 | MP Bauer et al., 2011 |
| LUMCMM19 0677 | pCD-METRO - | 106 | S | ECDIS 2013 | MP Bauer et al., 2011 |
| LUMCMM19 0678 | pCD-METRO - | 045 | S | ECDIS 2013 | MP Bauer et al., 2011 |
| LUMCMM19 0679 | pCD-METRO - | 014 | S | ECDIS 2013 | MP Bauer et al., 2011 |
| LUMCMM19 0680 | pCD-METRO - | 005 | S | ECDIS 2013 | MP Bauer et al., 2011 |
| LUMCMM19 0681 | pCD-METRO - | 014 | S | ECDIS 2013 | MP Bauer et al., 2011 |
| LUMCMM19 0682 | pCD-METRO - | 027 | S | ECDIS 2013 | MP Bauer et al., 2011 |
| LUMCMM19 0683 | pCD-METRO - | 476 | S | ECDIS 2013 | MP Bauer et al., 2011 |
| LUMCMM19 0684 | pCD-METRO - | 014 | S | ECDIS 2013 | MP Bauer et al., 2011 |
| LUMCMM19 0685 | pCD-METRO - | 027 | S | ECDIS 2013 | MP Bauer et al., 2011 |
| LUMCMM19 0686 | pCD-METRO - | 027 | S | ECDIS 2013 | MP Bauer et al., 2011 |
| LUMCMM19 0687 | pCD-METRO - | 018 | S | ECDIS 2013 | MP Bauer et al., 2011 |
| LUMCMM19 0688 | pCD-METRO - | 002 | S | ECDIS 2013 | MP Bauer et al., 2011 |
| LUMCMM19 0689 | pCD-METRO - | 045 | S | ECDIS 2013 | MP Bauer et al., 2011 |
| LUMCMM19 0690 | pCD-METRO - | 020 | S | ECDIS 2013 | MP Bauer et al., 2011 |
| LUMCMM19 0691 | pCD-METRO - | 027 | S | ECDIS 2013 | MP Bauer et al., 2011 |
| LUMCMM19 0692 | pCD-METRO - | 027 | S | ECDIS 2013 | MP Bauer et al., 2011 |
| LUMCMM19 0693 | pCD-METRO - | 078 | S | ECDIS 2013 | MP Bauer et al., 2011 |
| LUMCMM19 0694 | pCD-METRO - | 014 | S | ECDIS 2013 | MP Bauer et al., 2011 |
| LUMCMM19 0695 | pCD-METRO - | 027 | S | ECDIS 2013 | MP Bauer et al., 2011 |
| LUMCMM19 0696 | pCD-METRO - | 027 | S | ECDIS 2013 | MP Bauer et al., 2011 |
| LUMCMM19 0697 | pCD-METRO - | 081 | S | ECDIS 2013 | MP Bauer et al., 2011 |
| LUMCMM19 0698 | pCD-METRO - | 027 | S | ECDIS 2013 | MP Bauer et al., 2011 |
| LUMCMM19 0699 | pCD-METRO - | 027 | S | ECDIS 2013 | MP Bauer et al., 2011 |
| LUMCMM19 0700 | pCD-METRO - | 020 | S | ECDIS 2013 | MP Bauer et al., 2011 |
| LUMCMM19 0701 | pCD-METRO - | 045 | S | ECDIS 2013 | MP Bauer et al., 2011 |
| LUMCMM19 0702 | pCD-METRO - | 014 | S | ECDIS 2013 | MP Bauer et al., 2011 |
| LUMCMM19 0703 | pCD-METRO - | 014 | S | ECDIS 2013 | MP Bauer et al., 2011 |
| LUMCMM19 0704 | pCD-METRO - | 017 | S | ECDIS 2013 | MP Bauer et al., 2011 |
| LUMCMM19 0705 | pCD-METRO - | 057 | S | ECDIS 2013 | MP Bauer et al., 2011 |
| LUMCMM19 0706 | pCD-METRO - | 029 | S | ECDIS 2013 | MP Bauer et al., 2011 |
| LUMCMM19 0707 | pCD-METRO - | 014 | S | ECDIS 2013 | MP Bauer et al., 2011 |
| LUMCMM19 0708 | pCD-METRO - | 012 | S | ECDIS 2013 | MP Bauer et al., 2011 |
| LUMCMM19 0709 | pCD-METRO - | NT | S | ECDIS 2013 | MP Bauer et al., 2011 |
| LUMCMM19 0710 | pCD-METRO - | 010 | S | ECDIS 2013 | MP Bauer et al., 2011 |
| LUMCMM19 0711 | pCD-METRO - | 070 | S | ECDIS 2013 | MP Bauer et al., 2011 |
| LUMCMM19 0712 | pCD-METRO - | 020 | S | ECDIS 2013 | MP Bauer et al., 2011 |
| LUMCMM19 0713 | pCD-METRO - | 126 | S | ECDIS 2013 | MP Bauer et al., 2011 |
| LUMCMM19 0714 | pCD-METRO - | 053 | S | ECDIS 2013 | MP Bauer et al., 2011 |
| LUMCMM19 0715 | pCD-METRO - | 029 | S | ECDIS 2013 | MP Bauer et al., 2011 |
| LUMCMM19 0716 | pCD-METRO - | 081 | S | ECDIS 2013 | MP Bauer et al., 2011 |
| LUMCMM19 0717 | pCD-METRO - | 014 | S | ECDIS 2013 | MP Bauer et al., 2011 |
| LUMCMM19 0718 | pCD-METRO - | 014 | S | ECDIS 2013 | MP Bauer et al., 2011 |
| LUMCMM19 0719 | pCD-METRO - | 045 | S | ECDIS 2013 | MP Bauer et al., 2011 |
| LUMCMM19 0720 | pCD-METRO - | 014 | S | ECDIS 2013 | MP Bauer et al., 2011 |
| LUMCMM19 0721 | pCD-METRO - | 070 | S | ECDIS 2013 | MP Bauer et al., 2011 |
| LUMCMM19 0722 | pCD-METRO - | 027 | S | ECDIS 2013 | MP Bauer et al., 2011 |
| LUMCMM19 0723 | pCD-METRO - | 001 | S | ECDIS 2013 | MP Bauer et al., 2011 |
| LUMCMM19 0724 | pCD-METRO - | 001 | S | ECDIS 2013 | MP Bauer et al., 2011 |
| LUMCMM19 0725 | pCD-METRO - | 005 | S | ECDIS 2013 | MP Bauer et al., 2011 |
| LUMCMM19 0726 | pCD-METRO - | 001 | S | ECDIS 2013 | MP Bauer et al., 2011 |
| LUMCMM19 0727 | pCD-METRO - | 053 | S | ECDIS 2013 | MP Bauer et al., 2011 |
| LUMCMM19 0728 | pCD-METRO - | 114 | S | ECDIS 2013 | MP Bauer et al., 2011 |
| LUMCMM19 0729 | pCD-METRO - | 027 | S | ECDIS 2013 | MP Bauer et al., 2011 |
| LUMCMM19 0730 | pCD-METRO - | 014 | S | ECDIS 2013 | MP Bauer et al., 2011 |
| LUMCMM19 0731 | pCD-METRO - | 027 | S | ECDIS 2013 | MP Bauer et al., 2011 |
| LUMCMM19 0732 | pCD-METRO - | 024 | S | ECDIS 2013 | MP Bauer et al., 2011 |
| LUMCMM19 0733 | pCD-METRO - | 012 | S | ECDIS 2013 | MP Bauer et al., 2011 |
| LUMCMM19 0734 | pCD-METRO - | 027 | S | ECDIS 2013 | MP Bauer et al., 2011 |
| LUMCMM19 0735 | pCD-METRO - | 018 | S | ECDIS 2013 | MP Bauer et al., 2011 |
| LUMCMM19 0736 | pCD-METRO - | 014 | S | ECDIS 2013 | MP Bauer et al., 2011 |
| LUMCMM19 0737 | pCD-METRO - | 093 | S | ECDIS 2013 | MP Bauer et al., 2011 |
| LUMCMM19 0738 | pCD-METRO - | 027 | S | ECDIS 2013 | MP Bauer et al., 2011 |
| LUMCMM19 0739 | pCD-METRO - | 027 | S | ECDIS 2013 | MP Bauer et al., 2011 |
| LUMCMM19 0740 | pCD-METRO - | 027 | S | ECDIS 2013 | MP Bauer et al., 2011 |
| LUMCMM19 0741 | pCD-METRO - | 005 | S | ECDIS 2013 | MP Bauer et al., 2011 |
| LUMCMM19 0742 | pCD-METRO - | 020 | S | ECDIS 2013 | MP Bauer et al., 2011 |
| LUMCMM19 0743 | pCD-METRO - | 029 | S | ECDIS 2013 | MP Bauer et al., 2011 |
| LUMCMM19 0744 | pCD-METRO - | 027 | S | ECDIS 2013 | MP Bauer et al., 2011 |
| LUMCMM19 0745 | pCD-METRO - | 027 | S | ECDIS 2013 | MP Bauer et al., 2011 |
| LUMCMM19 0746 | pCD-METRO - | 027 | S | ECDIS 2013 | MP Bauer et al., 2011 |
| LUMCMM19 0747 | pCD-METRO - | 027 | S | ECDIS 2013 | MP Bauer et al., 2011 |
| LUMCMM19 0748 | pCD-METRO - | 005 | S | ECDIS 2013 | MP Bauer et al., 2011 |
| LUMCMM19 0749 | pCD-METRO - | 002 | S | ECDIS 2013 | MP Bauer et al., 2011 |
| LUMCMM19 0750 | pCD-METRO - | 005 | S | ECDIS 2013 | MP Bauer et al., 2011 |

| | | | | | |
|---|---|---|---|---|---|
| LUMCMM19 0751 | pCD-METRO - | 027 | S | ECDIS 2013 | MP Bauer et al., 2011 |
| LUMCMM19 0752 | pCD-METRO - | 027 | S | Tolevamer | TJ Louie et al., 2006 |
| LUMCMM19 0753 | pCD-METRO - | 019 | S | Tolevamer | TJ Louie et al., 2006 |
| LUMCMM19 0754 | pCD-METRO - | 139 | S | Tolevamer | TJ Louie et al., 2006 |
| LUMCMM19 0755 | pCD-METRO - | 027 | S | Tolevamer | TJ Louie et al., 2006 |
| LUMCMM19 0756 | pCD-METRO - | 075 | S | Tolevamer | TJ Louie et al., 2006 |
| LUMCMM19 0757 | pCD-METRO - | 001 | S | Tolevamer | TJ Louie et al., 2006 |
| LUMCMM19 0758 | pCD-METRO - | 001 | S | Tolevamer | TJ Louie et al., 2006 |
| LUMCMM19 0759 | pCD-METRO - | 053 | S | Tolevamer | TJ Louie et al., 2006 |
| LUMCMM19 0760 | pCD-METRO - | 003 | S | Tolevamer | TJ Louie et al., 2006 |
| LUMCMM19 0761 | pCD-METRO - | 002 | S | Tolevamer | TJ Louie et al., 2006 |
| LUMCMM19 0762 | pCD-METRO - | 570 | S | Tolevamer | TJ Louie et al., 2006 |
| LUMCMM19 0763 | pCD-METRO - | 056 | S | Tolevamer | TJ Louie et al., 2006 |
| LUMCMM19 0764 | pCD-METRO - | 001 | S | Tolevamer | TJ Louie et al., 2006 |
| LUMCMM19 0765 | pCD-METRO - | 001 | S | Tolevamer | TJ Louie et al., 2006 |
| LUMCMM19 0766 | pCD-METRO - | 001 | S | Tolevamer | TJ Louie et al., 2006 |
| LUMCMM19 0767 | pCD-METRO - | 027 | S | Tolevamer | TJ Louie et al., 2006 |
| LUMCMM19 0768 | pCD-METRO - | 001 | S | Tolevamer | TJ Louie et al., 2006 |
| LUMCMM19 0769 | pCD-METRO - | 176 | S | Tolevamer | TJ Louie et al., 2006 |
| LUMCMM19 0770 | pCD-METRO - | 001 | S | Tolevamer | TJ Louie et al., 2006 |
| LUMCMM19 0771 | pCD-METRO - | 001 | S | Tolevamer | TJ Louie et al., 2006 |
| LUMCMM19 0772 | pCD-METRO - | 808 | S | Tolevamer | TJ Louie et al., 2006 |
| LUMCMM19 0773 | pCD-METRO - | 027 | S | Tolevamer | TJ Louie et al., 2006 |
| LUMCMM19 0774 | pCD-METRO - | 019 | S | Tolevamer | TJ Louie et al., 2006 |
| LUMCMM19 0775 | pCD-METRO - | 075 | S | Tolevamer | TJ Louie et al., 2006 |
| LUMCMM19 0776 | pCD-METRO - | 080 | S | Tolevamer | TJ Louie et al., 2006 |
| LUMCMM19 0777 | pCD-METRO - | 014 | S | Tolevamer | TJ Louie et al., 2006 |
| LUMCMM19 0778 | pCD-METRO - | 009 | S | Tolevamer | TJ Louie et al., 2006 |
| LUMCMM19 0779 | pCD-METRO - | 027 | S | Tolevamer | TJ Louie et al., 2006 |
| LUMCMM19 0780 | pCD-METRO - | 027 | S | Tolevamer | TJ Louie et al., 2006 |
| LUMCMM19 0781 | pCD-METRO - | 027 | S | Tolevamer | TJ Louie et al., 2006 |
| LUMCMM19 0782 | pCD-METRO - | 014 | S | Tolevamer | TJ Louie et al., 2006 |
| LUMCMM19 0783 | pCD-METRO - | 002 | S | Tolevamer | TJ Louie et al., 2006 |
| LUMCMM19 0784 | pCD-METRO - | 023 | S | Tolevamer | TJ Louie et al., 2006 |
| LUMCMM19 0785 | pCD-METRO - | 001 | S | Tolevamer | TJ Louie et al., 2006 |
| LUMCMM19 0786 | pCD-METRO - | 126 | S | Tolevamer | TJ Louie et al., 2006 |
| LUMCMM19 0787 | pCD-METRO - | 014 | S | Tolevamer | TJ Louie et al., 2006 |
| LUMCMM19 0788 | pCD-METRO - | 001 | S | Tolevamer | TJ Louie et al., 2006 |
| LUMCMM19 0789 | pCD-METRO - | 005 | S | Tolevamer | TJ Louie et al., 2006 |
| LUMCMM19 0790 | pCD-METRO - | 016 | S | Tolevamer | TJ Louie et al., 2006 |
| LUMCMM19 0791 | pCD-METRO - | 014 | S | Tolevamer | TJ Louie et al., 2006 |
| LUMCMM19 0792 | pCD-METRO - | 036 | S | Tolevamer | TJ Louie et al., 2006 |
| LUMCMM19 0793 | pCD-METRO - | 023 | S | Tolevamer | TJ Louie et al., 2006 |
| LUMCMM19 0794 | pCD-METRO - | 010 | S | LUMC reference laboratory/RIVM | This study |
| LUMCMM19 0795 | pCD-METRO - | 010 | S | LUMC reference laboratory/RIVM | This study |
| LUMCMM19 0796 | pCD-METRO - | 010 | S | LUMC reference laboratory/RIVM | This study |
| LUMCMM19 0797 | pCD-METRO - | 010 | S | LUMC reference laboratory/RIVM | This study |
| LUMCMM19 0798 | pCD-METRO - | NT | S | LUMC reference laboratory/RIVM | This study |
| LUMCMM19 0799 | pCD-METRO - | 010 | S | LUMC reference laboratory/RIVM | This study |
| LUMCMM19 0800 | pCD-METRO - | 010 | S | LUMC reference laboratory/RIVM | This study |
| LUMCMM19 0801 | pCD-METRO - | 010 | S | LUMC reference laboratory/RIVM | This study |
| LUMCMM19 0802 | pCD-METRO - | 010 | S | LUMC reference laboratory/RIVM | This study |
| LUMCMM19 0803 | pCD-METRO - | 010 | S | LUMC reference laboratory/RIVM | This study |
| LUMCMM19 0804 | pCD-METRO - | 010 | S | LUMC reference laboratory/RIVM | This study |
| LUMCMM19 0805 | pCD-METRO - | 010 | S | LUMC reference laboratory/RIVM | This study |
| LUMCMM19 0806 | pCD-METRO - | 010 | S | LUMC reference laboratory/RIVM | This study |
| LUMCMM19 0807 | pCD-METRO - | 010 | S | LUMC reference laboratory/RIVM | This study |
| LUMCMM19 0808 | pCD-METRO - | NT | S | LUMC reference laboratory/RIVM | This study |
| LUMCMM19 0809 | pCD-METRO - | NT | S | LUMC reference laboratory/RIVM | This study |
| LUMCMM19 0810 | pCD-METRO - | 010 | S | LUMC reference laboratory/RIVM | This study |
| LUMCMM19 0811 | pCD-METRO - | 010 | S | LUMC reference laboratory/RIVM | This study |
| LUMCMM19 0812 | pCD-METRO - | 078 | S | LUMC reference laboratory/RIVM | This study |
| LUMCMM19 0813 | pCD-METRO - | 010 | S | LUMC reference laboratory/RIVM | This study |
| LUMCMM19 0814 | pCD-METRO - | 010 | S | LUMC reference laboratory/RIVM | This study |
| LUMCMM19 0815 | pCD-METRO - | 010 | S | LUMC reference laboratory/RIVM | This study |
| LUMCMM19 0816 | pCD-METRO - | 010 | S | LUMC reference laboratory/RIVM | This study |
| LUMCMM19 0817 | pCD-METRO - | 010 | S | LUMC reference laboratory/RIVM | This study |
| LUMCMM19 0818 | pCD-METRO - | 010 | S | LUMC reference laboratory/RIVM | This study |
| LUMCMM19 0819 | pCD-METRO - | 010 | S | LUMC reference laboratory/RIVM | This study |
| LUMCMM19 0820 | pCD-METRO - | 010 | S | LUMC reference laboratory/RIVM | This study |
| LUMCMM19 0821 | pCD-METRO - | 010 | S | LUMC reference laboratory/RIVM | This study |
| LUMCMM19 0822 | pCD-METRO - | NT | S | LUMC reference laboratory/RIVM | This study |
| LUMCMM19 0823 | pCD-METRO + | 010 | R | LUMC reference laboratory/RIVM | This study |
| LUMCMM19 0824 | pCD-METRO - | 010 | S | LUMC reference laboratory/RIVM | This study |
| LUMCMM19 0825 | pCD-METRO - | 010 | S | LUMC reference laboratory/RIVM | This study |
| LUMCMM19 0826 | pCD-METRO - | 010 | S | LUMC reference laboratory/RIVM | This study |
| LUMCMM19 0827 | pCD-METRO - | 010 | S | LUMC reference laboratory/RIVM | This study |
| LUMCMM19 0828 | pCD-METRO - | 010 | S | LUMC reference laboratory/RIVM | This study |
| LUMCMM19 0829 | pCD-METRO + | 010 | R | LUMC reference laboratory/RIVM | This study |
| LUMCMM19 0830 | pCD-METRO - | 010 | R | LUMC reference laboratory/RIVM | This study |
| LUMCMM19 0831 | pCD-METRO - | 010 | S | LUMC reference laboratory/RIVM | This study |
| LUMCMM19 0832 | pCD-METRO - | 010 | S | LUMC reference laboratory/RIVM | This study |
| LUMCMM19 0833 | pCD-METRO - | 010 | S | LUMC reference laboratory/RIVM | This study |
| LUMCMM19 0834 | pCD-METRO - | 010 | S | LUMC reference laboratory/RIVM | This study |
| LUMCMM19 0835 | pCD-METRO - | 010 | S | LUMC reference laboratory/RIVM | This study |
| LUMCMM19 0836 | pCD-METRO - | 010 | S | LUMC reference laboratory/RIVM | This study |
| LUMCMM19 0837 | pCD-METRO - | 010 | S | LUMC reference laboratory/RIVM | This study |
| LUMCMM19 0838 | pCD-METRO - | 010 | S | LUMC reference laboratory/RIVM | This study |
| LUMCMM19 0839 | pCD-METRO - | 010 | S | LUMC reference laboratory/RIVM | This study |
| LUMCMM19 0840 | pCD-METRO + | 010 | R | LUMC reference laboratory/RIVM | This study |
| LUMCMM19 0841 | pCD-METRO + | 010 | R | LUMC reference laboratory/RIVM | This study |
| LUMCMM19 0842 | pCD-METRO - | 010 | S | LUMC reference laboratory/RIVM | This study |
| LUMCMM19 0843 | pCD-METRO + | 010 | R | LUMC reference laboratory/RIVM | This study |
| LUMCMM19 0844 | pCD-METRO + | 010 | R | LUMC reference laboratory/RIVM | This study |
| LUMCMM19 0845 | pCD-METRO - | 010 | S | LUMC reference laboratory/RIVM | This study |
| LUMCMM19 0846 | pCD-METRO - | 010 | S | LUMC reference laboratory/RIVM | This study |
| LUMCMM19 0847 | pCD-METRO - | 010 | S | LUMC reference laboratory/RIVM | This study |
| LUMCMM19 0848 | pCD-METRO - | 078 | S | LUMC reference laboratory/RIVM | This study |
| LUMCMM19 0849 | pCD-METRO - | 013 | S | LUMC reference laboratory/RIVM | This study |
| LUMCMM19 0850 | pCD-METRO - | 198 | S | ECDC | This study |
| LUMCMM19 0851 | pCD-METRO - | 198 | S | ECDC | This study |
| LUMCMM19 0852 | pCD-METRO - | 198 | S | ECDC | This study |
| LUMCMM19 0853 | pCD-METRO - | 198 | S | ECDC | This study |
| LUMCMM19 0854 | pCD-METRO - | 198 | S | ECDC | This study |
| LUMCMM19 0855 | pCD-METRO - | 198 | S | ECDC | This study |
| LUMCMM19 0856 | pCD-METRO - | 198 | S | ECDC | This study |
| LUMCMM19 0857 | pCD-METRO - | 198 | S | ECDC | This study |
| LUMCMM19 0858 | pCD-METRO - | 198 | S | ECDC | This study |
| LUMCMM19 0859 | pCD-METRO - | 002 | S | ECDC | This study |
| LUMCMM19 0860 | pCD-METRO - | 198 | S | ECDC | This study |
| LUMCMM19 0861 | pCD-METRO - | 198 | S | ECDC | This study |
| LUMCMM19 0862 | pCD-METRO - | 198 | S | ECDC | This study |
| LUMCMM19 0863 | pCD-METRO - | 198 | S | ECDC | This study |
| LUMCMM19 0864 | pCD-METRO - | 198 | S | ECDC | This study |
| LUMCMM19 0865 | pCD-METRO - | 198 | S | ECDC | This study |
| LUMCMM19 0866 | pCD-METRO - | 198 | S | ECDC | This study |

| | | | | | |
|---|---|---|---|---|---|
| LUMCMM19 0867 | pCD-METRO - | 001 | S | ECDC | This study |
| LUMCMM19 0868 | pCD-METRO - | 027 | S | ECDC | This study |
| LUMCMM19 0869 | pCD-METRO - | 001 | S | ECDC | This study |
| LUMCMM19 0870 | pCD-METRO - | 001 | S | ECDC | This study |
| LUMCMM19 0871 | pCD-METRO - | 015 | S | ECDC | This study |
| LUMCMM19 0872 | pCD-METRO - | 001 | S | ECDC | This study |
| LUMCMM19 0873 | pCD-METRO - | 176 | S | ECDC | This study |
| LUMCMM19 0874 | pCD-METRO - | 001 | S | ECDC | This study |
| LUMCMM19 0875 | pCD-METRO - | 076 | S | ECDC | This study |
| LUMCMM19 0876 | pCD-METRO - | 001 | S | ECDC | This study |
| LUMCMM19 0877 | pCD-METRO - | 001 | S | ECDC | This study |
| LUMCMM19 0878 | pCD-METRO - | 001 | S | ECDC | This study |
| LUMCMM19 0879 | pCD-METRO - | 017 | S | ECDC | This study |
| LUMCMM19 0880 | pCD-METRO + | 010 | R | ECDC | This study |
| LUMCMM19 0881 | pCD-METRO - | 176 | S | ECDC | This study |
| LUMCMM19 0882 | pCD-METRO - | 027 | S | ECDC | This study |
| LUMCMM19 0883 | pCD-METRO - | 029 | S | ECDC | This study |
| LUMCMM19 0884 | pCD-METRO - | 176 | S | ECDC | This study |
| LUMCMM19 0885 | pCD-METRO - | 176 | S | ECDC | This study |
| LUMCMM19 0886 | pCD-METRO - | 027 | S | ECDC | This study |
| LUMCMM19 0887 | pCD-METRO - | 027 | S | ECDC | This study |
| LUMCMM19 0888 | pCD-METRO - | 027 | S | ECDC | This study |
| LUMCMM19 0889 | pCD-METRO - | 027 | S | ECDC | This study |
| LUMCMM19 0890 | pCD-METRO - | 027 | S | ECDC | This study |
| LUMCMM19 0891 | pCD-METRO - | 027 | S | ECDC | This study |
| LUMCMM19 0892 | pCD-METRO - | 010 | S | ECDC | This study |
| LUMCMM19 0893 | pCD-METRO - | NT | S | ECDC | This study |
| LUMCMM19 0894 | pCD-METRO - | 176 | S | ECDC | This study |
| LUMCMM19 0895 | pCD-METRO - | 216 | S | ECDC | This study |
| LUMCMM19 0896 | pCD-METRO - | 103 | S | ECDC | This study |
| LUMCMM19 0897 | pCD-METRO - | 003 | S | ECDC | This study |
| LUMCMM19 0898 | pCD-METRO - | 003 | S | ECDC | This study |
| LUMCMM19 0899 | pCD-METRO - | 003 | S | ECDC | This study |
| LUMCMM19 0900 | pCD-METRO - | 014 | S | ECDC | This study |
| LUMCMM19 0901 | pCD-METRO - | 001 | S | ECDC | This study |
| LUMCMM19 0902 | pCD-METRO - | 020 | S | ECDC | This study |
| LUMCMM19 0903 | pCD-METRO - | 001 | S | ECDC | This study |
| LUMCMM19 0904 | pCD-METRO - | 020 | S | ECDC | This study |
| LUMCMM19 0905 | pCD-METRO - | 003 | S | ECDC | This study |
| LUMCMM19 0906 | pCD-METRO - | 001 | S | ECDC | This study |
| LUMCMM19 0907 | pCD-METRO - | 001 | S | ECDC | This study |
| LUMCMM19 0908 | pCD-METRO - | 020 | S | ECDC | This study |
| LUMCMM19 0909 | pCD-METRO - | 001 | S | ECDC | This study |
| LUMCMM19 0910 | pCD-METRO - | 001 | S | ECDC | This study |
| LUMCMM19 0911 | pCD-METRO - | 010 | S | ECDC | This study |
| LUMCMM19 0912 | pCD-METRO - | 014 | S | ECDC | This study |
| LUMCMM19 0913 | pCD-METRO - | 001 | S | ECDC | This study |
| LUMCMM19 0914 | pCD-METRO - | 013 | S | ECDC | This study |
| LUMCMM19 0915 | pCD-METRO - | 001 | S | ECDC | This study |
| LUMCMM19 0916 | pCD-METRO - | 001 | S | ECDC | This study |
| LUMCMM19 0917 | pCD-METRO - | 014 | S | ECDC | This study |
| LUMCMM19 0918 | pCD-METRO - | 001 | S | ECDC | This study |
| LUMCMM19 0919 | pCD-METRO - | 014 | S | ECDC | This study |
| LUMCMM19 0920 | pCD-METRO - | 012 | S | ECDC | This study |
| LUMCMM19 0921 | pCD-METRO - | 643 | S | ECDC | This study |
| LUMCMM19 0922 | pCD-METRO - | 023 | S | ECDC | This study |
| LUMCMM19 0923 | pCD-METRO - | 012 | S | ECDC | This study |
| LUMCMM19 0924 | pCD-METRO - | 027 | S | MODIFY I+II | MH Wilcox et al., 2017 |
| LUMCMM19 0925 | pCD-METRO - | 027 | S | MODIFY I+II | MH Wilcox et al., 2017 |
| LUMCMM19 0926 | pCD-METRO - | 027 | S | MODIFY I+II | MH Wilcox et al., 2017 |
| LUMCMM19 0927 | pCD-METRO - | 027 | S | MODIFY I+II | MH Wilcox et al., 2017 |
| LUMCMM19 0928 | pCD-METRO - | 027 | S | MODIFY I+II | MH Wilcox et al., 2017 |
| LUMCMM19 0929 | pCD-METRO - | 027 | S | MODIFY I+II | MH Wilcox et al., 2017 |
| LUMCMM19 0930 | pCD-METRO - | 081 | S | MODIFY I+II | MH Wilcox et al., 2017 |
| LUMCMM19 0931 | pCD-METRO - | 027 | S | MODIFY I+II | MH Wilcox et al., 2017 |
| LUMCMM19 0932 | pCD-METRO - | 020 | S | MODIFY I+II | MH Wilcox et al., 2017 |
| LUMCMM19 0933 | pCD-METRO - | 017 | S | MODIFY I+II | MH Wilcox et al., 2017 |
| LUMCMM19 0934 | pCD-METRO - | 017 | S | MODIFY I+II | MH Wilcox et al., 2017 |
| LUMCMM19 0935 | pCD-METRO - | 001 | S | MODIFY I+II | MH Wilcox et al., 2017 |
| LUMCMM19 0936 | pCD-METRO - | 001 | S | MODIFY I+II | MH Wilcox et al., 2017 |
| LUMCMM19 0937 | pCD-METRO - | 001 | S | MODIFY I+II | MH Wilcox et al., 2017 |
| LUMCMM19 0938 | pCD-METRO - | 198 | S | MODIFY I+II | MH Wilcox et al., 2017 |
| LUMCMM19 0939 | pCD-METRO - | 027 | S | MODIFY I+II | MH Wilcox et al., 2017 |
| LUMCMM19 0940 | pCD-METRO - | 001 | S | MODIFY I+II | MH Wilcox et al., 2017 |
| LUMCMM19 0941 | pCD-METRO - | 001 | S | MODIFY I+II | MH Wilcox et al., 2017 |
| LUMCMM19 0942 | pCD-METRO - | 001 | S | MODIFY I+II | MH Wilcox et al., 2017 |
| LUMCMM19 0943 | pCD-METRO - | 027 | S | MODIFY I+II | MH Wilcox et al., 2017 |
| LUMCMM19 0944 | pCD-METRO - | 001 | S | MODIFY I+II | MH Wilcox et al., 2017 |
| LUMCMM19 0945 | pCD-METRO - | 027 | S | MODIFY I+II | MH Wilcox et al., 2017 |
| LUMCMM19 0946 | pCD-METRO - | 137 | S | MODIFY I+II | MH Wilcox et al., 2017 |
| LUMCMM19 0947 | pCD-METRO - | 027 | S | MODIFY I+II | MH Wilcox et al., 2017 |
| LUMCMM19 0948 | pCD-METRO - | 027 | S | MODIFY I+II | MH Wilcox et al., 2017 |
| LUMCMM19 0949 | pCD-METRO - | 231 | S | MODIFY I+II | MH Wilcox et al., 2017 |
| LUMCMM19 0950 | pCD-METRO - | 001 | S | MODIFY I+II | MH Wilcox et al., 2017 |
| LUMCMM19 0951 | pCD-METRO - | 027 | S | MODIFY I+II | MH Wilcox et al., 2017 |
| LUMCMM19 0952 | pCD-METRO - | 017 | S | MODIFY I+II | MH Wilcox et al., 2017 |
| LUMCMM19 0953 | pCD-METRO - | 001 | S | MODIFY I+II | MH Wilcox et al., 2017 |
| LUMCMM19 0954 | pCD-METRO - | 001 | S | MODIFY I+II | MH Wilcox et al., 2017 |
| LUMCMM19 0955 | pCD-METRO - | 176 | S | MODIFY I+II | MH Wilcox et al., 2017 |
| LUMCMM19 0956 | pCD-METRO - | 176 | S | MODIFY I+II | MH Wilcox et al., 2017 |
| LUMCMM19 0957 | pCD-METRO - | 176 | S | MODIFY I+II | MH Wilcox et al., 2017 |
| LUMCMM19 0958 | pCD-METRO - | 404 | S | MODIFY I+II | MH Wilcox et al., 2017 |
| LUMCMM19 0959 | pCD-METRO - | 027 | S | MODIFY I+II | MH Wilcox et al., 2017 |
| LUMCMM19 0960 | pCD-METRO + | 027 | R | MODIFY I+II | MH Wilcox et al., 2017 |
| LUMCMM19 0961 | pCD-METRO - | 176 | S | MODIFY I+II | MH Wilcox et al., 2017 |
| LUMCMM19 0962 | pCD-METRO - | 027 | S | MODIFY I+II | MH Wilcox et al., 2017 |
| LUMCMM19 0963 | pCD-METRO - | 176 | S | MODIFY I+II | MH Wilcox et al., 2017 |
| LUMCMM19 0964 | pCD-METRO - | 027 | S | MODIFY I+II | MH Wilcox et al., 2017 |
| LUMCMM19 0965 | pCD-METRO - | 027 | S | MODIFY I+II | MH Wilcox et al., 2017 |
| LUMCMM19 0966 | pCD-METRO - | 176 | S | MODIFY I+II | MH Wilcox et al., 2017 |
| LUMCMM19 0967 | pCD-METRO - | 027 | S | MODIFY I+II | MH Wilcox et al., 2017 |
| LUMCMM19 0968 | pCD-METRO - | 027 | S | MODIFY I+II | MH Wilcox et al., 2017 |
| LUMCMM19 0969 | pCD-METRO - | 027 | S | MODIFY I+II | MH Wilcox et al., 2017 |
| LUMCMM19 0970 (pu | pCD-METRO + | 010 | R | University General Hospital Gregorio Marañón, Spain | I Moura et al., 2013 |
| IB151 (published as P | pCD-METRO + | 010 | R | Institute for Medical Microbiology, Immunology and Hygiene, University of Cologne, Germany | E Piepenbrock et al., 2019 |

**Strains which were Whole Genome Sequenced**

| Name | Characteristics | PCR ribotype | MTZ resistance | Source | Reference |
|---|---|---|---|---|---|
| IB132 | pCD-METRO - | 020 | S | LUMC reference laboratory/RIVM | This study |
| IB133 | pCD-METRO + | 020 | R | LUMC reference laboratory/RIVM | This study |
| IB134 | pCD-METRO + | 020 | R | LUMC reference laboratory/RIVM | This study |
| IB135 | pCD-METRO + | 020 | R | LUMC reference laboratory/RIVM | This study |
| IB136 | pCD-METRO + | 020 | R | LUMC reference laboratory/RIVM | This study |
| IB137 | pCD-METRO - | 078 | S | LUMC reference laboratory/RIVM | This study |
| IB138 | pCD-METRO + | 010 | R | LUMC reference laboratory/RIVM | This study |

| | | | | | |
|---|---|---|---|---|---|
| IB139 | pCD-METRO - | 010 | S | LUMC reference laboratory/RIVM | This study |
| IB140 | pCD-METRO - | 010 | S | LUMC reference laboratory/RIVM | This study |
| IB141 | pCD-METRO - | 010 | S | LUMC reference laboratory/RIVM | This study |
| IB142 | pCD-METRO - | 010 | S | LUMC reference laboratory/RIVM | This study |
| IB143 | pCD-METRO + | 010 | R | LUMC reference laboratory/RIVM | This study |
| IB144 | pCD-METRO + | 010 | R | LUMC reference laboratory/RIVM | This study |
| IB145 | pCD-METRO + | 010 | R | LUMC reference laboratory/RIVM | This study |
| IB146 | pCD-METRO + | 010 | R | LUMC reference laboratory/RIVM | This study |
| IB147 | pCD-METRO + | 010 | R | LUMC reference laboratory/RIVM | This study |
| IB148 | pCD-METRO + | 010 | R | LUMC reference laboratory/RIVM | This study |
| IB149 | pCD-METRO + | 010 | R | LUMC reference laboratory/RIVM | This study |

**Other strains used in this study**

| Name | Characteristics | PCR ribotype | MTZ resistance | Source | Reference |
|---|---|---|---|---|---|
| AP38 | *C. difficile* 630Δ*erm* pRPF185; Thia$^R$ | NA | S | LUMC | AM Oliveira Paiva et al., 2016 |
| CA434 | *E. coli* HB101 [F$^-$ mcrB mrr hsdS20($r_B^- m_B^-$) recA14 leuB6 ara-14 proA2 lacY1 galK2 xyl-5 mtl-1 rspL20(Sm$^R$)glnV44λ$^-$] with plasmid R702 | NA | R | Lab stock | D. Purdy et al., 2002 |
| DH5α | *E. coli* F– endA1 glnV44 thi-1 recA1 relA1 gyrA96 deoR nupG purB20 φ80dlacZΔM15 Δ(lacZYA-argF)U169, hsdR17(rK–mK+), λ– | NA | R | Lab stock | Laboratory stock |
| MDS42 | *E. coli* MG1655 multiple-deletion strain 72 ΔdinB ΔpolB ΔumuDC 120 ΔIS609 ΔpatD ΔydcV ΔydcU ΔydcT ΔydcS ΔydcR ΔhicA ΔhicB ΔyncJ ΔydcP ΔydcN ΔydcO ΔydcM ΔrecA(1819) | NA | R | Lab stock | Scarab Genomics, LLC |
| EVE17 | *C. difficile* 630Δerm; ErmS; RifR | 012 | S | LUMC | This study |
| IB30 | *C. difficile* 630Δ*erm* pIB20; Thia$^R$ | 012 | S | LUMC | This study |
| IB82 | *C. difficile* 630Δerm pIB76; ThiaR | 012 | S | LUMC | This study |
| IB83 | *C. difficile* 630Δerm pIB77; ThiaR | 012 | S | LUMC | This study |
| IB84 | *C. difficile* 630Δerm pIB78; ThiaR | 012 | S | LUMC | This study |
| IB89 | *C. difficile* 630Δerm pIB79; ThiaR | 012 | S | LUMC | This study |
| IB90 | *C. difficile* 630Δ*erm* pIB80; Thia$^R$ | 012 | S | LUMC | This study |
| IB125 | *C. difficile* 630Δ*erm* pIB86; Thia$^R$ | 012 | R | LUMC | This study |
| WKS1710 | *C. difficile* 630Δerm tcda::CT tcdB::CT; ErmR | 012 | S | LUMC | This study |

**Supplementary table 1: all strains used in this study**

| Name | Relevant features | Source/reference |
|---|---|---|
| pRPF185 | tetR $P_{tet}$-s*luc* $^{opt}$; catP | A Oliveira Paiva et al., 2016 |
| pCD-METRO | (see: accession number ERZ807316) | This study |
| pIB20 | pRPF185 $P_{cd0716}$-s*luc* $^{opt}$; catP | This study |
| pIB76 | tetR Ptet-nimB; catP | This study |
| pIB77 | tetR Ptet-metallohydrolase; catP | This study |
| pIB78 | Pnat-metallohydrolase-nimB; catP | This study |
| pIB79 | tetR Ptet-metallohydrolase-nimB; catP | This study |
| pIB80 | tetR $P_{tet}$-*gusA -pCD-METROreplicon* | This study |
| pIB86 | pCD-METRO containing shuttle fragment; pCD-METRO-*oriT-pBR322-traJ catP*; catP | This study |

**Supplementary table 2: plasmids used in this study**

| Name | Sequence (5’->3’) | Description | Source |
|---|---|---|---|
| oBH-1 | CCTCGTAGAATCCGGTGCAA | Forward primer annealing to *orf6* of pCD-METRO | This study |
| oBH-2 | TATTTCCTTGCCGCTGAGGT | Reverse primer annealing to *orf6* of pCD-METRO | This study |
| oBH-3 | GCAGAGCGTTGTGGTATTCG | Forward primer annealing to *orf6* of pCD-METRO | This study |
| oBH-4 | GTATTTCCTTGCCGCTGAGG | Reverse primer annealing to *orf6* of PCD-METRO | This study |
| oBH-5 | AGTAGGCGGTCGGTACTTCT | Forward primer annealing to *orf5* of pCD-METRO | This study |
| oBH-6 | CTTCCGTTCCGCTTGTTTGG | Reverse primer annealing to *orf5* of pCD-METRO | This study |
| oBH-7 | GGCTCAGACACTTCTACCGC | Forward primer annealing to *orf7* of pCD-METRO | This study |
| oBH-8 | CCCCTTCCAGGGTGTTTTCT | Reverse primer annealing to *orf7* of pCD-METRO | This study |
| oBH-9 | GGGCTATAAGACCGACTGGC | Forward primer annealing to *orf2* of pCD-METRO | This study |
| oBH-10 | AACGGTCTCTACCTCCGTCA | Reverse primer annealing to *orf2* of pCD-METRO | This study |
| oBH-11 | ACTTACACTGCAAACGGTGC | Forward primer annealing to *orf8* of pCD-METRO | This study |
| oBH-12 | TCGTTGCTTGTGAGGTGAGT | Reverse primer annealing to *orf8* of pCD-METRO | This study |
| oIB-103 | CATATAAGATAAAAAATATATTGAAAAATAC | Forward primer used for sequencing *rpoB* | This study |
| oIB-104 | GATATGACCTAGCAAAAGTTGGTAG | Forward primer used for sequencing *rpoB* | This study |
| oIB-105 | AGAATCACCATATAGAAAATTTGATAAAG | Forward primer used for sequencing *rpoB* | This study |
| oIB-106 | GGAGAAACTGAACTTACTGCTGAG | Forward primer used for sequencing *rpoB* | This study |
| oIB-107 | GACAGATGAAGACCAAGAAATAGAAG | Forward primer used for sequencing *rpoB* | This study |
| oIB-108 | ATTGTCTTATTTTTTCTGGAGAAGCC | Forward primer used for sequencing *rpoB* | This study |
| oIB-120 | GCCGAAAAAGAAAAACTGCCGGG | Sequencing primer for pCD-METRO$^{shuttle}$ | This study |
| oIB-121 | CTTACTCAAACGGAAGTGATGAGAG | Sequencing primer for pCD-METRO$^{shuttle}$ | This study |
| oIB-122 | GCGTTTATAGTACACTGGCTTGTG | Sequencing primer for pCD-METRO$^{shuttle}$ | This study |
| RP314 | GAAGGTTGACCACGGTATCAT | Forward qPCR primer for *catR* | EM Ransom et al., 2015 |
| RP315 | CGCAACGGTATGGAAACAATC | Reverse qPCR primer for *catR* | EM Ransom et al., 2015 |
| TEQ009 | AAGAGCTGGATTCGAAGTGCGTGA | Forward qpCR primer for *rpoB* | EM Ransom et al., 2015 |
| TEQ010 | ACCGATATTTGGTCCCTCTGGAGT | Reverse qPCR primer for *rpoB* | EM Ransom et al., 2015 |
| oWKS-1241 | CACCGACGAGCAAGGCAAGACCG | Sequencing primer for pCD-METRO$^{shuttle}$ | E Van Eijk et al., 2019 |
| oWKS-1387 | CAGATGAGGGCAAGCGGATG | Sequencing primer for pCD-METRO$^{shuttle}$ | This study |
| oWKS-1388 | CGTCGGTGAGCCAGAGTTTC | Sequencing primer for pCD-METRO$^{shuttle}$ | This study |
| oWKS-1537 | TAGGGTAACAAAAAACACCG | Sequencing primer for pCD-METRO$^{shuttle}$ | PT van Leeuwen et al., 2016 |
| oWKS-1539 | GGATTTCACATTTGCCGTTTTGTAAAC | Sequencing primer for pCD-METRO$^{shuttle}$ | PT van Leeuwen et al., 2016 |
| oWKS-1540 | GATCTTTTCTACGGGGTCTGAC | Sequencing primer for pCD-METRO$^{shuttle}$ | PT van Leeuwen et al., 2016 |
| oWKS-1574 | AAACAAACCACCGCTGGTAG | Sequencing primer for pCD-METRO$^{shuttle}$ | This study |
| oWKS-1656 | TAGCGGATCCAAGCGTTCTGAACGCACTG | Sequencing primer for pCD-METRO$^{shuttle}$ | This study |
| oWKS-1658 | TAGCGGATCCGGGCTTACTTCTGGGTATCC | Sequencing primer for pCD-METRO$^{shuttle}$ | This study |
| oWKS-1659 | TAGCGGTACCTGTTGCCTGCTTCCTGTATG | Sequencing primer for pCD-METRO$^{shuttle}$ | This study |
| oWKS-1661 | TAGCGGTACCATCCACGCACCACAACAC | Sequencing primer for pCD-METRO$^{shuttle}$ | This study |
| oWKS-1663 | CAAGACGGTCGGCGTTGCGCTCGAAGATGGATAAAATAAAATAGAGGCTATAAATAGC | Forward primer for PCR of shuttle fragment *pBR322ori-catP-oriT-traJ* | This study |
| oWKS-1664 | CGATAAATCTTGATTTGATGAAGTACAAGGTTAGTAGGTGCTTTTTTAAAAC | Reverse primer for PCR of shuttle fragment *pBR322ori-catP-oriT-traJ* | This study |
| oWKS-1678 | CCATCTTCGAGCGCAACG | Sequencing primer for pCD-METRO$^{shuttle}$ | This study |

**Supplementary table 3: oligonucleotides used in this study**
